# Connectome Similarity Varies with Global Metastability and Movie Content

**DOI:** 10.64898/2025.12.14.694198

**Authors:** Tiago Duarte-Pereira, André M Cravo, Claudinei E Biazoli

**Affiliations:** Center of Mathematics, Computing and Cognition. Federal University of ABC, São Bernardo do Campo, São Paulo, Brazil; National Institute of Science and Technology on Social and Affective Neuroscience, INCT-SANI, São Paulo, Brazil

**Keywords:** Functional MRI, Brain Dynamics, Functional Connectivity, Naturalistic Stimuli

## Abstract

**Introduction:** Recent evidence supports that non-linear segments of brain activity carries most of the individual-specific information which allows unique identification based on functional connectomes (e.g. *connectome fingerprints)* in MEG. In fMRI, metastability has emerged as a reliable and robust summary measure of brain dynamics. Here, we set out to test the hypothesis that metastability is associated with connectome fingerprinting in naturalistic fMRI. We further examined whether this putative association varies with time scales, across brain functional networks, and depending on the movie content.

**Methods:** We analyzed fMRI data from 86 participants of the *Naturalistic Neuroimaging Database*. Data was collected during movie-watching of 10 entire movies of different genres. Global and network-level metastability were calculated using an fMRI-appropriate proxy measure based on the variability of spatial coherence. The within-subject stability (*Iself*) and between-subjects similarity (*Iothers*) of individual functional connectomes were also calculated. Linear mixed-effects models and appropriate contrasts were used to test associations between metastability and fingerprinting indices.

**Results:** At the whole-brain level and considering data from all movies, an association across windows was found between metastability, *Iself*, and *Iothers*, with stronger effects observed for larger windows, such as the 600-second window (*Iself*: *b* = −0.13, SE = 0.032, *p* < 0.001; *I*others: *b* = −0.23, SE = 0.028, *p* < 0.001). Associations were also observed with distinct magnitudes within the eight large-scale functional networks, with particularly strong effects in the Motor Network (*I*self: *b* = −0.34, SE = 0.027, *p* < 0.001; *I*others: *b* = −0.27, SE = 0.021, *p* < 0.001). Although no whole-brain differences were observed, movie content exhibited distinct association patterns within networks, notably for the prediction of *Iself* in the Visual Network for *500 Days of Summer* (Δ = −0.22, SE = 0.11, *p* = 0.043). For *Iothers*, significant between-movie differences were found in the AMN (Δ = −0.25, SE = 0.062, *p* < 0.001), Dorsal Attention (Δ = 0.16, SE = 0.08, *p* = 0.038), Ventral Attention (Δ = 0.14, SE = 0.060, *p* = 0.017), and Visual Networks (Δ = −0.17, SE = 0.081, *p* = 0.031).

**Conclusion:** Metastability is systematically related to functional connectome fingerprints. This relationship is time-scale dependent, varies across functional networks, and is sensitive to movie content. Our findings bridge nonlinear brain dynamics and connectome identifiability, highlighting naturalistic paradigms as a powerful framework for individualized precision neuroimaging.

## 1 INTRODUCTION

Computational Neuroscience has been achieving new and important results by modeling brain function as a dynamical system (Cabral et al., 2022; Fraiman et al., 2009; Herzog et al., 2023). Recently, signatures of metastability of the nervous system have emerged as a reliable summary measure of brain dynamics based on dynamical systems modeling approaches (Hancock et al., 2023). Metastability is a property of some non-linear systems that oscillates between transient stable and unstable states, and can be measured using fMRI (Hancock et al., 2024). Metastability has been shown to vary with distinct levels of and repertoire of consciousness states (Cavanna et al., 2018), including anesthesia and during psychedelic states (Hutchison et al., 2014; Tagliazucchi et al., 2014). Particularly, metastability was shown to vary during movie-watching. (Kringelbach et al., 2023).

In a parallel line of investigation, the variability of individual functional connectomes has been consistently shown to allow unique identification (i.e., connectome fingerprinting) across neuroimaging techniques, including fMRI (Finn et al., 2015), fNIRS (De Souza Rodrigues et al., 2019) and MEG (Da Silva Castanheira et al., 2021). A recent study using MEG (Sorrentino et al., 2023) has shown that short nonlinear patterns of activation explain most of the individual variability leading to unique identification. In fMRI, individual identification was shown to oscillate through time (Ville et al., 2021). In light of these recent findings supporting nonlinear dynamics as idiosyncratic, we hypothesize that global metastability predicts the individual functional connectome in fMRI.

Both metastability and connectome fingerprints also vary differentially across large-scale functional networks (Amico et al., 2014; Meer et al., 2020; Wens et al., 2019) and during development (Vanderwal et al., 2021). These network-level idiosyncrasies might be important for understanding the mechanisms underlying the connectome uniqueness and identifiability (Larabi et al., 2021). Moreover, clarifying the potential relation between brain dynamics and connectome fingerprint have many potential applications in social neuroscience (Parkinson et al., 2018), learning (Jangraw et al., 2023) and clinical settings (Fan et al., 2020; Sorrentino et al., 2021).

Finally, we hypothesize that the relationship between metastability and connectome fingerprints during movie-watching is stimuli-sensitive. For that, we took advantage of the acclaimed naturalistic turn in neuroimaging (Ibanez et al., 2024; Sonkusare et al., 2019). Naturalistic stimuli such as movies, music, and narratives provide rich cognitive and affective situations (Saarimäki, 2021), aids in reducing head motion artifact, increasing arousal (Frew et al., 2022), and consequently reducing sleep during acquisition (Tagliazucchi and Laufs, 2014). Indeed, naturalistic fMRI have shown to yield results that are more reliable compared to traditional resting-state protocols (Wang et al., 2017). However, much current research in the field employs short films (Nguyen et al., 2016; Rieck et al., 2020) or only excerpts from larger movies (Lahnakoski et al., 2017; Raz et al., 2012). Here we additionally intend to highlight the importance of considering the ecological nature of the stimuli by using data acquired during watching the original content and full-length movies.

Hence, we used the Naturalistic Neuroimaging Dataset (NNDb, Aliko et al., 2020), comprising fMRI data from 86 volunteers watching one of 10 selected films (minimum 91-minutes) to test our hypotheses regarding the relation between metastability and connectome fingerprints. To further investigate this relation, we decomposed connectome fingerprint in two indices: *Iself,* indicating within-subject stability along time*; Iother,* indicating subject-to-others similarity (Amico and Goñi, 2018). Considering previous MEG findings linking greater identifiability to nonlinear dynamics, we expected to observe association between metastability and the idiosyncratic component of the signal (i.e., *Iself*). We also expected to find stronger associations in primary sensorimotor networks compared to heteromodal networks (Vanderwal et al., 2021). Finally, we expected that associations between metastability and fingerprinting should vary with movie stimulus content (e.g., specific movie genre).

## 2 METHODS

### 2.1 Participants

The Naturalistic Neuroimaging Dataset (NNDb) comprises 86 participants (42 females, 18–58 years, M = 26.81, SD = 10.09 years) scanned while watching one of 10 different films spanning 10 different movie genres. The participants were all right-handed, with normal or lens-corrected vision, and had never watched the film before. Detailed demographics for each subject are described in the original paper (Aliko et al., 2020).

### 2.2 Stimuli

The stimuli consisted of 10 entire films spanning 10 different genres (Table 1), featuring diverse narratives and varying durations (the shortest film lasting 91-minutes and the longest lasting 154-minutes). Due to technical limitations inherent in fMRI, breaks (2-6 breaks) were necessary every 40-50 minutes (93.94% of breaks occurred every 41.54 minutes (SD = 10.47) during data collection. These breaks were carefully timed to respect the narrative flow and minimize interference. Additionally, some participants experienced additional pauses due to mild or severe drowsiness, technical adjustments (volume, headphones, cleaning of glasses lenses), and technical malfunctions. These issues were addressed, filtered, or excluded to maintain data quality.

**Table 1:** Description of the stimuli from the *NNDb*.

| MOVIE | GENRE | SUBJECTS | DURATION<br>(min.) | TRs |
| --- | --- | --- | --- | --- |
| 500 Days of Summer (2009) | Romance | 20 (10 females,<br>$M=27.7$,<br>$SD=10.14$) | 91 | 5470 |
| Citizenfour (2014) | Documentary | 18 (9 females,<br>$M=27$, $SD=10.51$) | 113 | 6804 |
| The Usual Suspects (1995) | Crime | 6 (3 females, M=23.17, SD=2.23) | 106 | 6102 |
| Pulp Fiction (1994) | Action | 6 (3 females, M=22.67, SD=4.50) | 154 | 8882 |
| The Shawshank Redemption (1994) | Drama | 6 (3 females, M=22.17, SD=4.17) | 142 | 8181 |
| The Prestige (2006) | Thriller | 6 (3 females, M=37, SD=16.42) | 130 | 7515 |
| Back to the Future (1985) | Sci-fi | 6 (3 females, M=22.17, SD=2.48) | 116 | 6674 |
| Split (2016) | Horror | 6 (3 females, M=22.67, SD=2.50) | 117 | 6739 |
| Little Miss Sunshine (2006) | Comedy | 6 (3 females, M=35.67, SD=14.72) | 101 | 5900 |
| 12 Years A Slave (2013) | Historical | 6 (3 females, M=27.17, SD=12.09) | 134 | 7715 |

We used the full data of all 86 subjects to investigate the putative relationship between measures independently of the narrative. In order to investigate the effects of the movie genres, we used the two movies with more subjects: 500 Days of Summer (20 subjects) and Citizenfour (18 subjects). The first one is a non-conventional romantic comedy movie with a scrambled chronology and a fast pacing. The second movie is a documentary depicting extensive interviews and dialogues between the director, a reporter and the former NSA analyst Edward Snowden. In comparison to the romantic comedy, the documentary has a much slower pace, a linear narrative and more silent moments.

### 2.3 fMRI Acquisition

All scans (anatomical and functional) were run in a 1.5 T Siemens MAGNETOM Avanto with a 32 channel head coil (Siemens Healthcare, Erlangen, Germany). A multiband EPI was used (TR = 1 s, TE = 54.8 ms, flip angle of 75°, 40 interleaved slices, resolution = 3.2 mm isotropic), with 4x multiband factor and no in-plane acceleration. To enable a reasonable comfort and quality of visualization of the film, a mirror reversing LCD projector was used to generate a rear-projection screen measuring 22.5 cm × 42 cm with a field of view angle of 19.0°. Non-magnetic Sensimetrics S14 scanner earphones were used for immersion and noise-reducing.

The display equipment was automated alongside the scanner, ensuring that movie breaks also paused the equipment, facilitating concatenation of breaks to guarantee correct synchronization throughout.

### 2.4 fMRI Preprocessing and Atlases

A brief description of the preprocessed pipeline (Aliko et al., 2020) to fMRI data includes: slice-timing correction, despiking, volume registration by aligning the timepoints to the mean functional image of the centre timeseries, MNI alignment, 6mm full-width smoothing, detrending with six demeaned motion regressors, and timing correction.

We used three different atlases to define cortical areas: the Glasser’s (Glasser et al., 2016), Gordon’s (Gordon et al., 2017) and the Harvard-Oxford atlas (Maldjian et al., 2003), with 360, 333 and 133 parcels, respectively (see Supplementary Material). Following consistency checks and removal of atlas-related artifacts, analyses were conducted using the Gordon atlas as it proved more suitable. We used 8 distinct large-scale functional networks: Somatomotor (SMN, 46 ROIs); Default Mode (DMN, 41 ROIs); Cíngulo-Opercular which corresponds to the Action-Mode (Dosenbach et al., 2024; AMN, 40 ROIs); Visual (VIN, 39 ROIs); Dorsal Attentional (DAN, 32 ROIs); Auditory (AUN, 24 ROIs); Frontoparietal (FPN, 24 ROIs); and Ventral Attentional (VAN, 23 ROIs).

### 2.5 Functional Connectivity inflation artifact at long acquisitions

Recently, a study by Korponay et al. (2024) raised the issue of systemic low-frequency oscillations (sLFO), an physiological noise artifact related to arousal during long fMRI acquisitions, which standard preprocessing methods were not addressing and leading to inflated Functional Connectivity values. It was extensively demonstrated through distinct scans specifications and 4 different cohorts: MIC, HCP, PSU and YMRRC. The Regressor Interpolation at Progressive Time Delays (RIPTiDe), implemented by the *rapidtide* software package, was proposed as a solution. This denoising is based on filtering the LFO band (0.009–0.15 Hz) and using it to extract, refine and regress from the LFO-filtered global mean BOLD signal. A detailed full description can be found in previous studies (Tong et al., 2019). We conducted an analysis to check whether the same occurred in our data (Results in Supplementary Material).

### 2.6 Measurements

To measure fMRI signals metastability, we used the standard deviation of spatial coherence as a proxy for each subject. This is the most suitable measure for fMRI data (Hellyer et al., 2014; Hancock et al., 2024), considering its lower temporal resolution compared to other neuroimaging techniques (i.e., EEG, fNIRS, MEG).

For identifiability measures, we defined a fingerprinting matrix using Pearson correlation and vectorization (Finn et al., 2015). We further decomposed the fingerprinting matrix in two complementary identifiability metrics, *Iself* and *Iothers* (Amico and Goñi, 2018). The idiosyncratic portion of functional connectivity is embedded in the diagonal of the fingerprinting matrix while shared connectivity patterns can be captured by the non-diagonal elements. *Iself* thus corresponds to the numerical value of the correlation of a person’s signal with their own signal at a different time. *Iothers* correspond to the mean value of all the correlations between a person’s signal and the rest of the subject’s signal.

These fingerprinting matrix indices have received different labels in the literature, including: (1) Stability and Similarity of the connectomes (Vanderwal et al., 2021), (2) Intra and Inter Subject-Variability (Amico & Goñi, 2018); and (3) *Iself* and *Iothers* indices. To avoid confusion with the use of metastability in this work, we adopted the terminology proposed by Amico and Goni (2018), and refer to these measures as *Iself* (for the diagonal elements of the fingerprinting matrix) and *Iother*s (for the average of the off-diagonal elements).

To take advantage of the dataset length and content richness, the measures were calculated with a sliding window approach, allowing to capture the metrics dynamics across the data, and to test different window sizes. To cover a wide range of information inside the sliding window, we run the computations based on a 60, 300 and 600-seconds window-size. Whole-brain and large-scale functional networks values of *Iself* and *Iothers* were accessed dividing the sliding window and using the halves to compute each fingerprinting matrix through the data, while metastability was calculated by the entire sliding window length.

**Figure 1:**
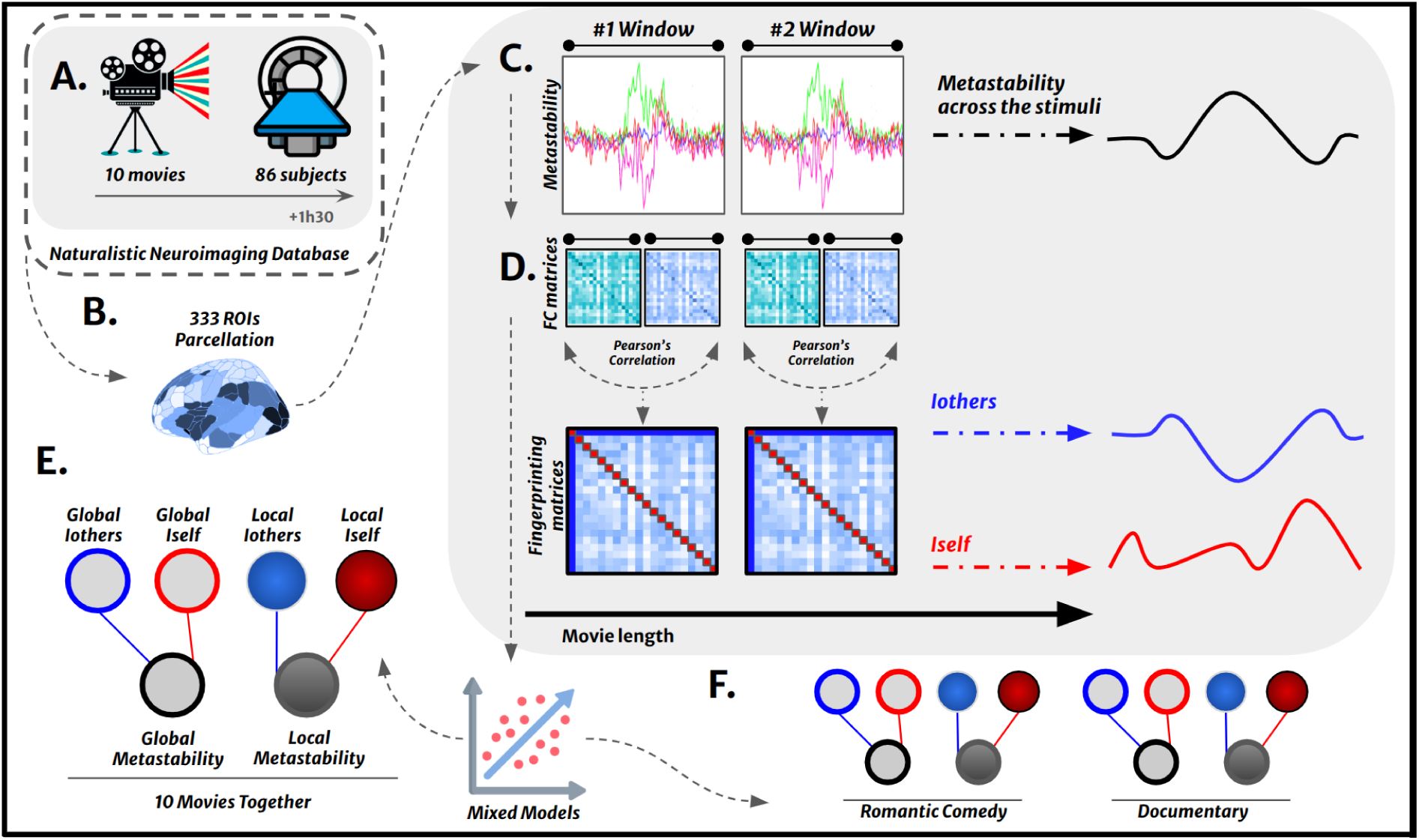
Study Pipeline. **A:** We use the *Naturalistic Neuroimaging Database (NNDb),* consisting of 86 subjects who watched an entire movie while in the MRI scanner. Each person watched one of 10 different films, which range from 91 to 154 minutes (see Table 1). **B:** The signal from all participants underwent dimensionality reduction using the 333 ROIs Gordon parcellation. **C:** Spatial coherence was calculated as the absolute value of the euclidean distance between the value of the region and the mean of the regions, while metastability was defined as the standard deviation of the specified time interval. **D:** The fingerprinting matrix was calculated as the Pearson correlation between subjects FC matrices in the same dataset. For each fingerprinting matrix, we calculated two measures:: the *Iself*, which is the value of comparing their FC matrix with themselves (Iself = <a_ii_>), and the *Iothers*, which is the average comparison of their FC matrix with all other subjects (Iothers = <a_ij_>) **E:** We fitted four distinct models: (i) All Movies models for whole-brain (333 ROIs), with metastability predicting each one of the fingerprinting indexes; (ii) All Movies model for each index and each one of the 8 large-scale functional networks; **F:** (iii) Whole-brain comparative models between the Romantic Comedy (*500 Days of Summer*) and the Documentary (*Citizenfour*) for both indexes; (iv) Network-level models comparing the two movies for both indexes. All models included age, sex, windows, movie genre and motion parameters as covariates.

### 2.7 Statistical Analyses

All statistical analyses were performed in RStudio 4.4.0. First we tested the hypothesis of a relationship between metastability and each fingerprinting index (*Iself* and *Iothers*). We fit a GMMs (i.e.: general mixed models) for each one of the dependent variables, *Iself* and *Iothers,* with metastability as the main predictor, Subjects included as random effects and controlling for each *window* and *movie genre* and for *age*, *sex* and six *motion parameters* for each subject. We employed the *lme4* package which uses the restricted maximum likelihood (REML) to estimate the model parameters. A recent paper brought light again on an old discussion about models complexity and reduction in interpretability (Rohrer and Arel-Bundock, 2025). Based on that, we employed the *marginaleffects* and *emmeans* packages, which allows us to make sense of the coefficients appropriately based on estimated quantities. The target quantity, or estimand, of this model will be the slope of the prediction curve for each dependent variable.For testing the relationship between fingerprinting indexes and metastability within the large-scale networks, the same model features were maintained, adding the categorical variable *Networks* as a covariate. Finally, to explore the potential effect of stimulus content in the relation between metastability and fingerprinting, at whole-brain and large-scale networks scale. For this, we computed average comparisons between the variable *movie genres*. As aforementioned, considering differences in the narrative structure and also the number of subjects, this analysis will focus on two movies: *500 Days of Summer* (N=20) and *Citizenfour* (N=18).

## 3 RESULTS

### 3.1 - Fingerprinting and global metastability across movies and timescales

First, we calculated descriptive statistics of the main metrics (metastability, *Iself* and *Iothers*) across all movies and windows. Results for the 60-second and 300-seconds time-windows and alternative parcelations are shown in Supplementary Material. In the main manuscript, we describe the results for 600-seconds time-windows in Gordon parcelation.

*Pulp Fiction* presented the lower Global Metastability index (M=0.08, SE=0.003, CI[0.057 0.118]) and *The Prestige* presented the higher (M=0.11, SE=0.004, CI[0.079 0.144]). The two movies of interest, *500 Days of Summer* and *Citizenfour*, presented similar values (M=0.10, SE=0.003, CI[0.062 0.141]; M=0.10, SE=0.002, CI[0.072 0.133]). For *Iself* and *Iothers*, *Usual Suspects* presented the lower *Iself* (M=0.64, SE=0.006, CI[0.595 0.0693]) and *The Prestige* presenting the higher *Iself* (M=0.70, SE=0.005, CI[0.658 0.748]); and *500 Days of Summer* (M=0.65, SE=0.004, CI[0.600 0.718]) and Citizenfour (M=0.67, SE=0.004, CI[0.616 0.736]) presented intermediate values. For *Iothers*, the lower and the higher values were respectively for *Back to the Future* (M=0.37, SE=0.002, CI[0.351 0.395]) and *Split* (M=0.43, SE=0.002, CI[0.411 0.450]); with similar values for *500 Days of Summer* (M=0.38, SE=0.001, CI[0.357 0.405]) as for *Citizenfour* (M=0.38, SE=0.001. CI[0.361 0.408]) (Figure 2). The full table of results for all movies can be found in Supplementary Material.

**Figure 2:**
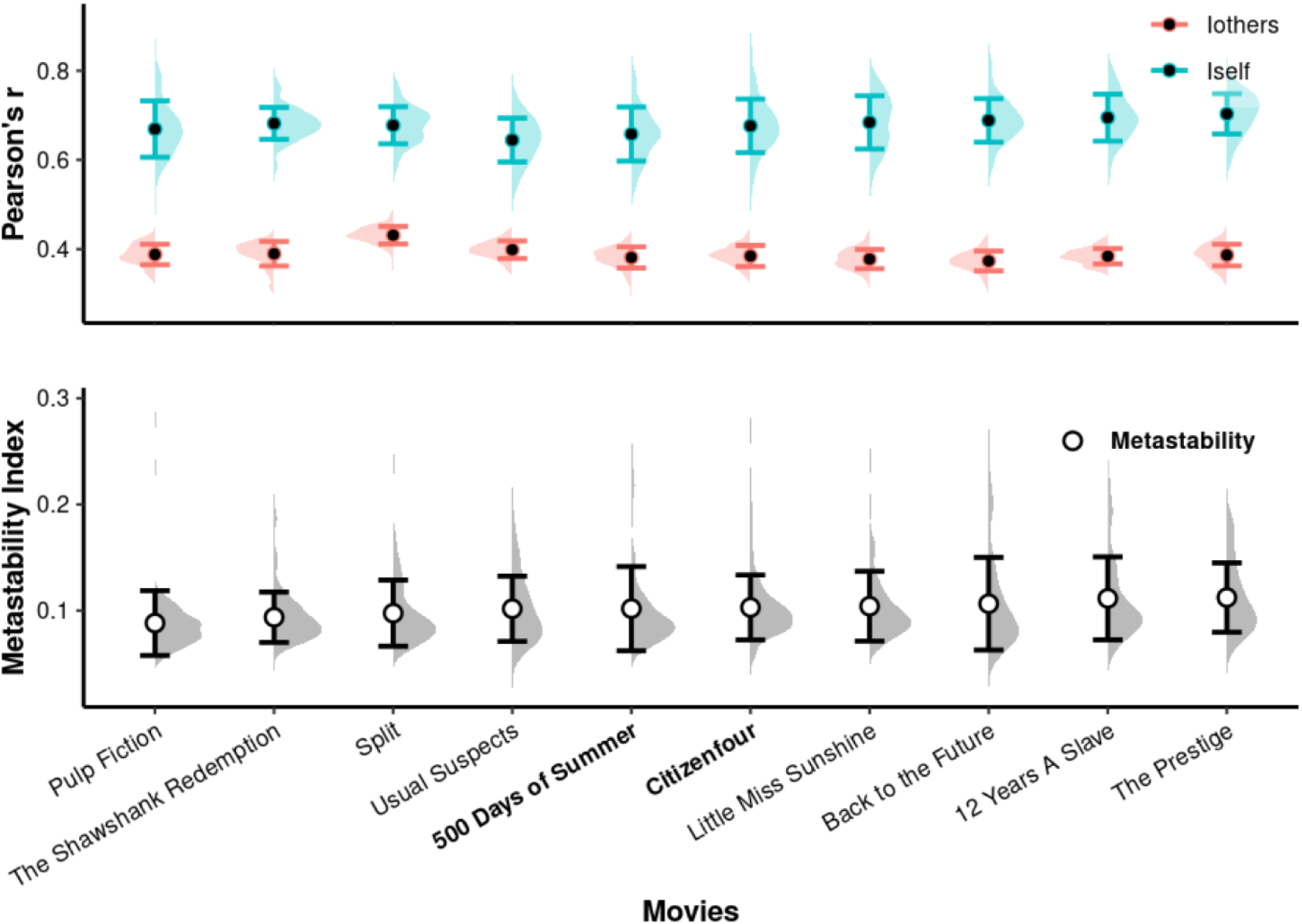
Descriptive statistics of *Iself, Iothers* and Metastability measures across movies for 600-seconds sliding windows. All the 86 subjects from the NNDb dataset were included. For further and sensitivity analyses, the 2 movies with most subjects were considered (500 Days of Summer, N=20; and Citizenfour, N=18). All the other 8 movies have the same amount of subjects (N=6). **Top:** Distribution, mean and standard deviation of *Iself* and *Iothers* measures. *Iself* represents a measure of the within-subject connectome stability and is calculated as the correlation between two same-subject functional connectivity matrices, or the diagonal part of the functional connectome fingerprinting matrix. *Iothers* represents a measure of between-subjects connectome similarity and is calculated as the correlation between a subject’s functional connectivity matrix and the average functional connectivity matrix of all other subjects, or the average of the non-diagonal part of the fingerprinting matrix. The plot shows the 95% confidence intervals and violin plots across movies of *Iself* (in blue) and *Iothers* (in red) measured as Pearson’s correlation r index, presented for *Iself* ranging values between 0.64 to 0.70, and for *Iothers* 0.37 to 0.43, of correlation. **Bottom:** Distribution, mean and standard deviation of Metastability, a measure derived from complex systems modelling that considers that the brain works at a regime that alternates between moments of integration and segregation. For fMRI, due to low time resolution, a proxy measure of this regime is reached by calculating the standard deviation of the spatial coherence. Metastability values are dimensionless, and reflect variability rather than a physical quantity, see Methods for a more detailed description.

Considering only the 2 movies of interest, the same above descriptive statistics were calculated across three windows of distinct sizes (60-seconds, 300-seconds and 600-seconds). *Iothers* and *Iself* presented a increase in it values with the increase of the window size, meanwhile *Metastability* does not presented visible differences. For the 60-seconds window size, 500 *Days of Summer* presented similar values compared to Citizenfour of *Iself* (M=0.24, SE=0.001, CI[0.176 0.310]; M=0.24, SE=0.001, CI[0.173 0.320]), I*others* (M=0.11, SE=0.0003, CI[0.098 0.128]; M=0.11, SE=0.0003, CI[0.100 0.128]) and *Metastability* (M=0.09, SE=0.001, CI[0.040 0.136]; M=0.09, SE=0.0009, CI[0.048 0.129]). For the 300-seconds window size, both movies presented respectively similarities in the three measures (*Iself:* M=0.53, SE=0.003, CI[0.466 0.601]; M=0.54, SE=0.003, CI[0.475 0.615]) | *Iothers:* M=0.29, SE=0.001, CI[0.274 0.318]; M=0.30, SE=0.001, CI[0.274 0.321] | *Metastability*: M=0.10, SE=0.002, CI[0.057 0.141]; M=0.10, SE=0.001, CI[0.068 0.133]). 600-seconds values were described in the plot above.

**Figure 3:**
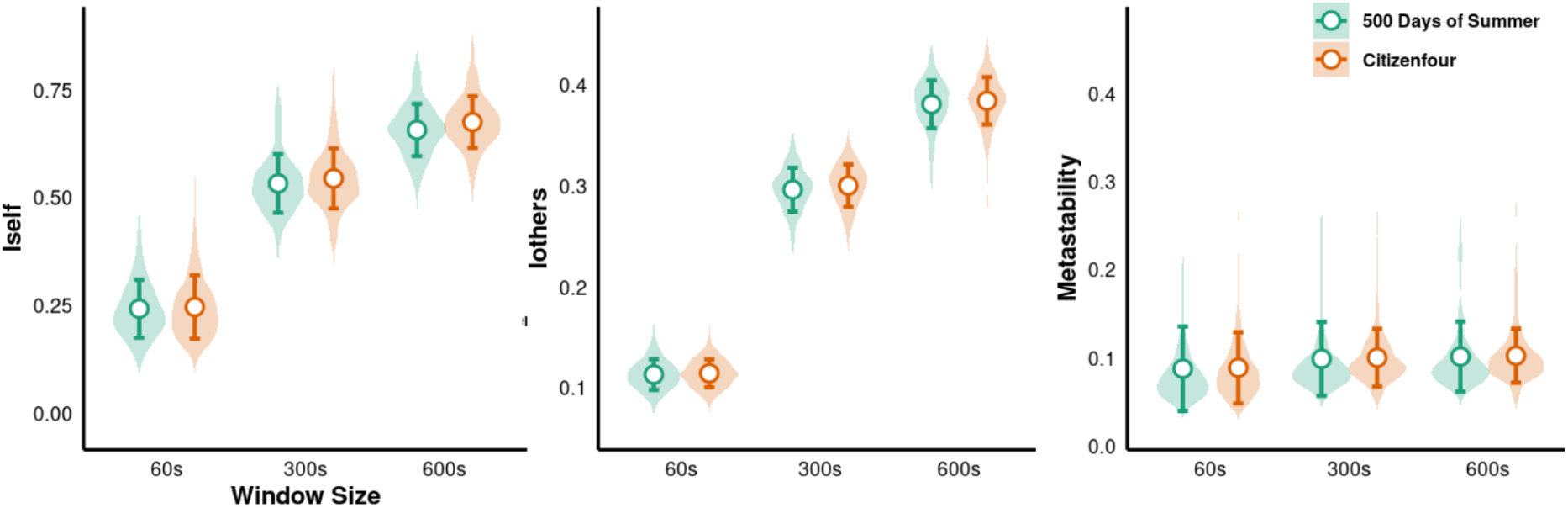
Violin plots with 95% confidence intervals of the three measures: *Iself* (left), *Iothers* (middle) and *Metastability* (right). Were calculated for 2 movies, *500 Days of Summer* (in green) and *Citizenfour* (in orange), and for three sliding windows: 60-seconds, 300-seconds and 600-seconds size.

In addition, we calculated for both movies the three measures across the movie data for the 600-seconds window size (for the 60-seconds and 300-seconds window size, see Supplementary Material). The two movies have distinct lengths, *500 Days of Summer* is 91-minutes long while *Citizenfour* is 113-minutes long, this is reflected in the amount of windows for each movie, which for this case are 9 as 11 windows respectively and it is indicated in the plot below. For *Iself*, with the peak of correlation in *500 Days of Summer* (M=0.66, SE=0.013, CI[0.655 0.681]) was in the 3rd window (between 20-minutes and 30-minutes of the movie) and the minimum value (M=0.64, SE=0.015, CI[0.633 0.664]) was in 6th window (between 50-minutes and 60-minutes); for *Citizenfour*, the peak (M=0.69, SE=0.015, CI[0.681 0.712]) was in the 8th window (between 1h10 and 1h20 of the movie) and the minimum value (M=0.66, SE=0.015, CI[0.651 0.682]) was in the following window (between 1h20 and 1h30 of the movie). For *Iothers, 500 Days of Summer* presenting peak (M=0.38, SE=0.005, CI[0.380 0.391]) at the 5th window and minimum value (M=0.37, SE=0.005, CI[0.370 0.382]) at 2nd window; at the same time, *Citizenfour* presented peak (M=0.38, SE=0.005, CI[0.383 0.394]) at the 3rd window and minimum (M=0.38, SE=0.004, CI[0.376 0.385]) at the 1st window. For *Metastability*, *500 Days of Summer* showed a peak in the 2nd window (M=0.11, SE=0.010, CI[0.095 0.117]) and minimum in the 7th window (M=0.09, SE=0.008, CI[0.090 0.106]) while *Citizenfour* showed a peak in the 9th window (M=0.11, SE=0.011, CI[0.101 0.123]) and a minimum value in the the 6th window (M=0.09, SE=0.004, CI[0.087 0.095]).

**Figure 4:**
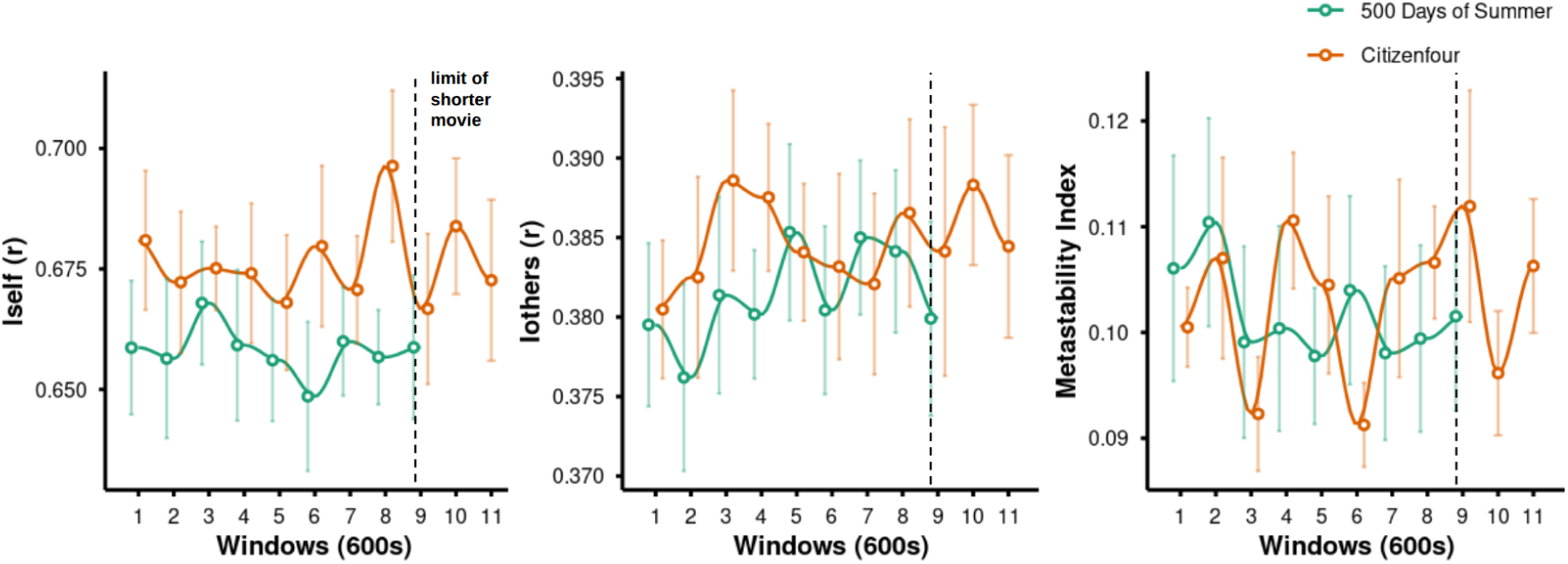
Line plot with 95% confidence intervals of the three measures: *Iself* (left), *Iothers* (middle) and *Metastability* (right) across the two movies (*500 Days of Summer* in green and *Citizenfour* in orange) data in a 600-seconds sliding window size. Distinct movie lengths are reflected with a distinct amount of time points, 9 and 11, respectively, with a dotted line describing the limit of the shorter movie, in this case, 500 Days of Summer (91-minutes long).

### 3.2 - Connectome identifiability is associated with global metastability

As expected, an association was found between both connectome fingerprinting indexes and global metastability (for all movies, N=86 and 333 ROIs), even after controlling for age, sex, movie genre, time-window and movement parameters. These results are robust across different window durations.

For 60-seconds sliding windows, *Iself* presented a significant positive association with metastability (b=0.08, SE=0.010, p<0.001), while no significant association was observed for *Iothers* (p=0.286). For the 300-seconds window, a negative association with metastability was observed for both *Iself* (b=-0.05, SE=0.023, p<0.016) and *Iothers* (b=-0.20, SE=0.020, p<0.001). The same pattern of association was observed for the 600-seconds window (*Iself -* b=-0.13, SE=0.032, p<0.001; *Iothers* - b=-0.23, SE=0.028, p<0.001).

**Figure 8.**
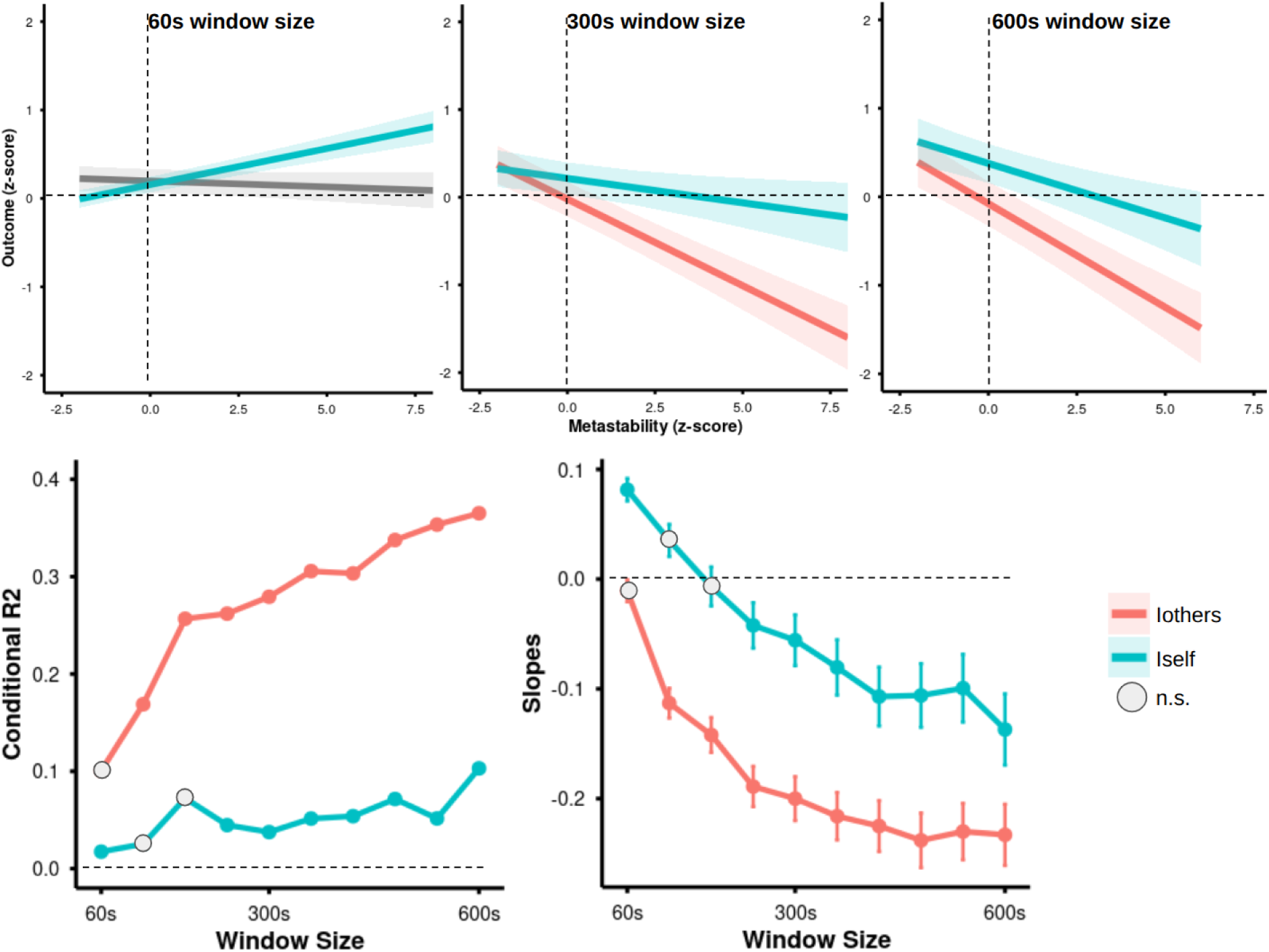
Results of the association between *Iself* and *Iothers* with global metastability after controlling for age, sex, movie Genre, window position and movement. Results of the association presented as regression slopes for three window sizes of (60-seconds, 300-seconds and 600-seconds). *Iself* (in blue) is positively associated with metastability for 60-seconds time-windows and negatively associated with metastability for time-windows greater than 240-second. For *Iothers,* a consistent negative association with metastability was observed for time-windows longer than 60-seconds window. As expected, conditional R^2^ increases with window duration, with higher values for *Iothers* for all windows.

### 3.3 - Fingerprinting and metastability associations in large-scale networks

Associations between functional connectome fingerprinting indexes and metastability were also observed at the large-scale networks level and for all the 8 networks investigated (AMN, Auditory, DMN, Dorsal Attentional, FPN, Motor, Ventral Attentional and Visual). Here, we present the results for the 600-second time window.

Significant negative associations were observed for both fingerprinting measures and metastability for all networks, except for *Iothers* in Ventral Attentional Network. Results for prediction of both measures within networks are summarized below (Table 2).

**Table 2:** Results for prediction of *Iself* and *Iothers* by metastability within networks for ‘All Movies’ model.

| Networks (*All Movies*) | *Isel* | *Ioth*ers |
| --- | --- | --- |
| AMN | b=-0.20, SE=0.027, p<0.001 | b=-0.17, SE=0.021, p<0.001 |
| Auditory | b=-0.21, SE=0.028, p<0.001 | b=-0.20, SE=0.022, p<0.001 |
| DMN | b=-0.22, SE=0.031, p<0.001 | b=-0.11, SE=0.024, p<0.001 |
| Dorsal Attentional | b=-0.22, SE=0.034, p<0.001 | b=-0.09, SE=0.027, p=0.001 |
| FPN | b=-0.18, SE=0.026, p<0.001 | b=-0.05, SE=0.020, p=0.007 |
| Motor | b=-0.34, SE=0.027, p<0.001 | b=-0.27, SE=0.021, p<0.001 |
| Ventral Attentional | b=-0.22, SE=0.027, p<0.001 | b=-0.02, SE=0.021, p=0.210 |
| Visual | b=-0.29, SE=0.034, p<0.001 | b=-0.15, SE=0.026, p<0.001 |

**Figure 9.**
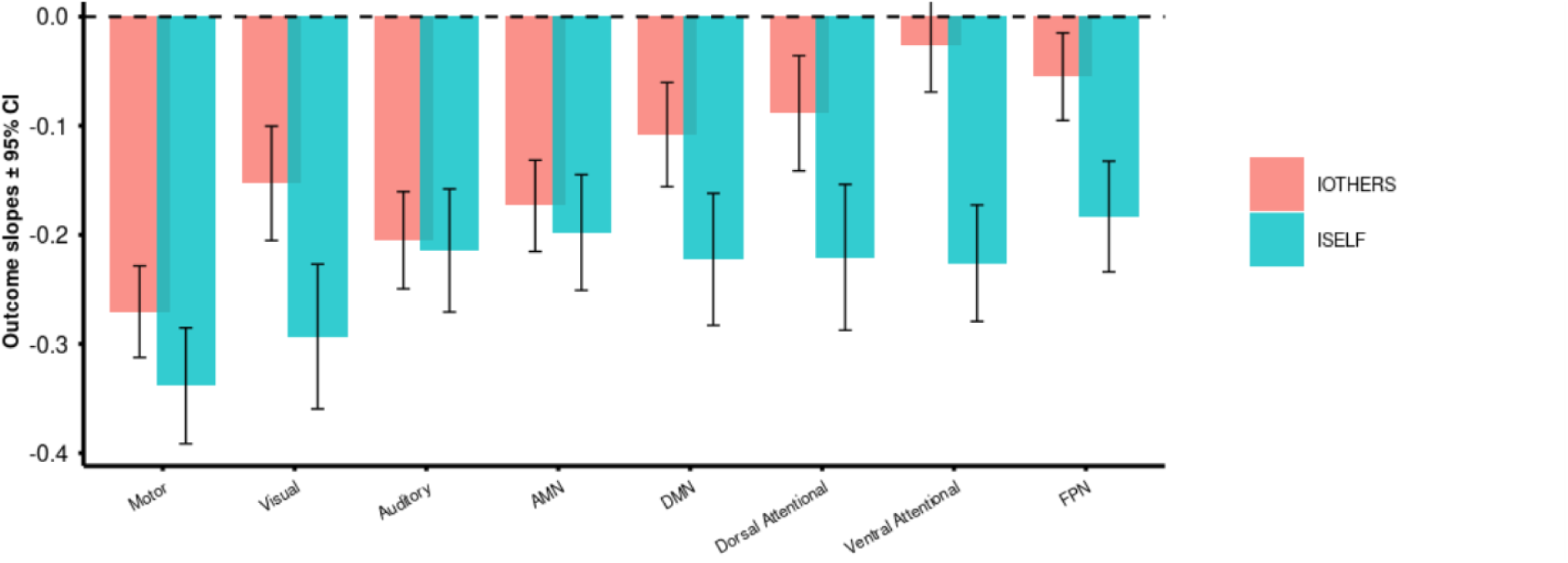
Network level associations between *Iself* and *Iothers* with metastability for 8 large-scale networks (Action Mode Network - AMN; Auditory Network; Default Mode Network - DMN; Dorsal Attentional Network; Fronto-Parietal Network - FPN; Motor Network; Ventral Attentional Network; and Visual Network), controlling for age, sex, movie Genre, windows and movement. Results of the analysis using 600-seconds windows and all 10 movies data (N=86). **Top:** Association estimates with 95% confidence intervals, with *Iself* results in blue and *Iothers* in red. Significant negative association with metastability were observed for all except Ventral Attentional Networks for the *Iothers* model.

### 3.4 - Association between identifiability and metastability related to stimulus content

*500 Days of Summer* (N=20) is a high paced and non-cronological romantic comedy while *Citizenfour* (N=18) is a slow paced and sober tone documentary about former NSA analyst Edward Snowden. Both movies presented in the descriptive statistics similarities in the three measures of interest independently on the window. We interrogated our models regarding the movie watched by the subjects focusing on these two movies based on the sample size.

At whole-brain level, a significant and quite similar negative association between *Iothers* and metastability was observed for both movies (*500 Days of Summer*: b=-0.37, SE=0.060, p<0.001 | *Citizenfour*: b=-0.36, SE=0.074, p<0.001). The same pattern was observed for the relation between *Iself* and global metastability (*500 Days of Summer*: b=-0.23, SE=0.063, p<0.001 | *Citizenfour*: b=-0.29, SE=0.078, p<0.001). At the large-scale networks level, a significant difference between movie-related slopes of the association between *ISelf* and global metastability was observed specifically in the Visual Network (Δ=-0.22, SE=0.11, p=0.043). Results for prediction within all networks are summarized on the table below (Table 3).

**Table 3:** Results for prediction of *Iself* by metastability within networks.

| Networks (*Isel*) | *500 Days of Summer* | *Citizenfour* |
| --- | --- | --- |
| AMN | $b=-0.28$, $SE=0.057$, $p<0.001$ | $b=-0.27$, $SE=0.066$, $p<0.001$ |
| Auditory | $b=-0.26$, $SE=0.051$, $p<0.001$ | $b=-0.32$, $SE=0.074$, $p<0.001$ |
| DMN | $b=-0.26$, $SE=0.067$, $p=0.001$ | $b=-0.39$, $SE=0.072$, $p<0.001$ |
| Dorsal Attentional | $b=-0.21$, $SE=0.066$, $p=0.001$ | $b=-0.39$, $SE=0.093$, $p<0.001$ |
| FPN | $b=-0.16$, $SE=0.048$, $p=0.001$ | $b=-0.27$, $SE=0.072$, $p<0.001$ |
| Motor | $b=-0.35$, $SE=0.058$, $p<0.001$ | $b=-0.52$, $SE=0.067$, $p<0.001$ |
| Ventral Attentional | $b=-0.21$, $SE=0.050$, $p<0.001$ | $b=-0.29$, $SE=0.067$, $p<0.001$ |
| Visual | $b=-0.43$, $SE=0.062$, $p<0.001$ | $b=-0.20$, $SE=0.094$, $p=0.035$ |

Regarding the relation between *Iothers* and metastability across the movies, a significant value was found for *500 Days of Summer* (b=-0.25, SE=0.040, p<0.001) but not for *Citizenfour* (b=-0.00, SE=0.047, p=0.887) in Action Mode Network. Calculating the contrast between networks slopes, we indeed found significant differences for AMN (Δ=-0.25, SE=0.062, p<0.001), Dorsal Attentional (Δ=0.16, SE=0.08, p=0.038), Ventral Attentional (Δ=0.14, SE=0.060, p=0.017) and Visual Networks (Δ=-0.17, SE=0.081, p=0.031). Results for prediction within all networks are summarized on the table below (Table 4).

**Table 4:** Results for prediction of *Iothers* by metastability within networks.

| Networks (*Iothers*) | *500 Days of Summer* | *Citizenfour* |
| --- | --- | --- |
| AMN | b=-0.25, SE=0.040, p<0.001 | b=-0.00, SE=0.047, p=0.887 |
| Auditory | b=-0.26, SE=0.037, p<0.001 | b=-0.29, SE=0.052, p<0.001 |
| DMN | b=-0.13, SE=0.048, p=0.005 | b=-0.21, SE=0.051, p<0.001 |
| Dorsal Attentional | b=-0.06, SE=0.047, p=0.145 | b=-0.23, SE=0.066, p=0.004 |
| FPN | b=-0.00, SE=0.034, p=0.816 | b=-0.11, SE=0.051, p=0.022 |
| Motor | b=-0.26, SE=0.041, p<0.001 | b=-0.28, SE=0.047, p<0.001 |
| Ventral Attentional | b=-0.02, SE=0.041, p=0.444 | b=-0.11, SE=0.048, p=0.017 |
| Visual | b=-0.19, SE=0.045, p<0.001 | b=-0.02, SE=0.067, p=0.760 |

**Figure 10.**
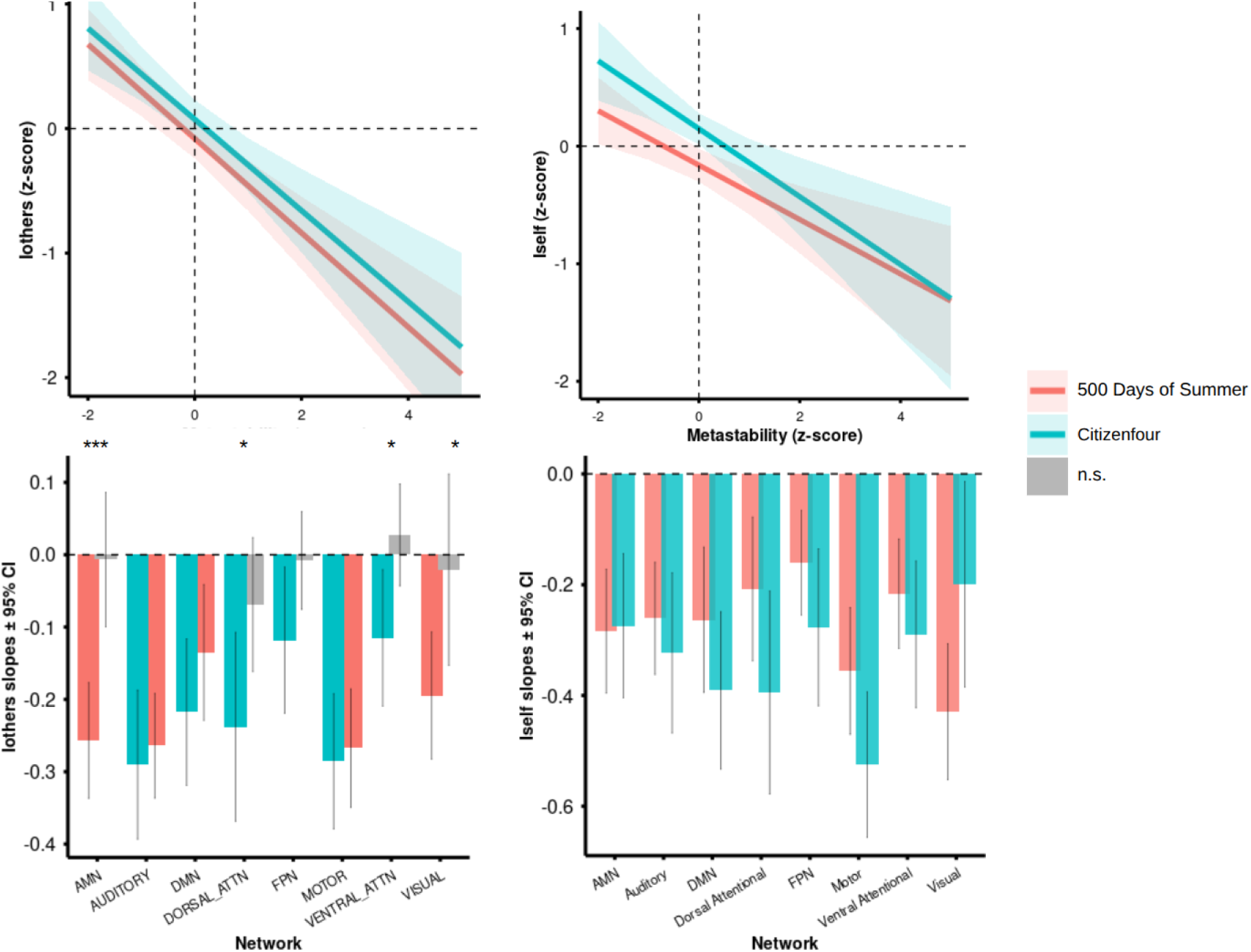
Associations of *Iself* and *Iothers* with *Metastability* at the whole-brain and across 8 large-scale networks separately for *500 Days of Summer* (N=20, in red) and *Citizenfour* (N=18, in blue). Results of models controlling for age, sex, windows and movement and 600-seconds windows. **Top:** Slope curves for *Iself* and *Iothers* relations with metastability for both movies present a quite similar negative association. **Bottom Left:** Results for *Iothers* associations with metastability for both movies and at the networks level. Statistically significant differences between the slopes were observed in Action Mode, Visual, Dorsal and Ventral Attentional Networks. present highlighted statistical differences. FPN, despite no statistical difference, also is highlighted in *Citizenfour*. **Bottom Right:** No statistical difference between network slopes for the different movies were observed regarding the association between *Iself* and metastability at the network level.

## 4 DISCUSSION

Both fingerprinting analyses and metastability measures are promising approaches for moving towards individualized precision neuroimaging. In line with previous evidence in MEG (Sorrentino et al., 2023) supporting that nonlinear functional dynamics (which can be reliably summarized by metastability) underlie functional connectome identifiability, we found a strong and consistent association between metastability and fingerprinting indexes in fMRI. Moreover, this association was shown to vary for different time-scales, across functional networks and with naturalistic stimuli content.

In our dataset, metastability and both fingerprinting indexes were generally similar across all movies. Interestingly, the *Iothers* values for the suspense/horror movie *Split* (2016) was considerably higher compared to all other movies. Previous work had suggested that suspense movies are indeed related with higher intersubject correlation values (Hasson et al., 2008), particularly for acute rather than sustained fear (Hudson et al., 2020).

A positive association between *Iself* (*i.e.*, stability within-subjects) and metastability was found but exclusively in shorter window sizes (*i.e.*, 60-seconds). These results seem to be more directly in line with those observed in MEG (Sorrentino et al., 2023). For longer time-windows, however, the inverse relation was consistently observed. The association between global metastability and *Iothers* (similarity between-subjects), on the other hand, remained consistently negative across time scales.

We speculate that the negative associations for both fingerprinting indexes with metastability for larger time-windows may be due to: (i) brain functional dynamics at these scales might be driven by sustained higher-order stimulus contents that make identifying patterns to vary over time: so, a time-window with higher metastability would lead to both a lower *Iself* and *Iothers*; or (ii) the time-windows used are larger than both idiosyncratic and shared functional features, which might have a lower duration, and our models are capturing slower affectively and cognitively relevant fluctuations; or (iii) our models are capturing slower artefactual fluctuations. Though this third possibility can not be completely ruled out, our further analyses do not support the results being driven by systemic low-frequency fluctuations (Supplementary Material). In short, long length movie-watching may lead individuals’ brains to maintain their idiosyncrasy while manifesting some shared functioning features which drives cognition.

The investigated films encompass quite different narratives styles: *500 days of Summer* is a fast paced, full of events non-cronological narrative, with a high visual-attentional demand or *film tyranny* (Loschky et al, 2015); *Citizenfour*, on the contrary, is a slow-paced, full of details and dense in information interview with former NSA analyst Edward Snowden, a narrative demanding sustained attention. Interestingly, we found that the relations between fingerprinting and metastability varied with movies in visual and attentional networks. A particularly strong difference was observed for the *Iothers* and metastability relation in the recently proposed Action Mode Network (Dosenbach et al., 2024).

It is important to highlight some limitations of the present investigation. fMRI data, especially for longer durations, is highly susceptible to motion artifacts that can have confounded our results, even in the high quality NNDb dataset and using all standard denoising procedures and including motion as a covariate in all of our models. We conducted sensitivity analyses with and without motion correction to also test the possibility of the corrections inducing yet another artifact. Those sensitivity analyses results were consistent with the main results present here but replication of our findings in other datasets is warranted. Scanner heating can also induce artifacts in long-duration scans. We indirectly tested the impact of this trend by analyzing data that was filtered for FC inflation (see Supplementary Material). Additionally, our results for the specific movies have a relatively small sample size and expanding these results for larger datasets and other movies would be necessary to confirm the specific patterns of the association between metastability and fingerprinting. Furthermore, though the currently most reliable measure of fMIR nonlinear dynamics, the proxy metastability can be tested and extended using proper whole-brain modelling.

Previous studies with event segmentation, anticipation of segmentation and incorporation of quantitative and qualitative features of the movie (Baldassano et al., 2017; Lee et al., 2021; Cohen et al., 2022) are promising ways to explore the association between metastability and fingerprinting. Finally, our results and a proper naturalistic account of movies as narrative experiences, could foster personalized neuroimaging approaches.

## 6 SUPPLEMENTARY MATERIAL

Supplementary Materials - Paper 1

## 7 DATA AND CODE AVAILABILITY

https://github.com/TiagoDuaarte/metastability-fingerprinting-naturalistic

## 8 AUTHOR CONTRIBUTIONS

Tiago Duarte-Pereira conceived the study, designed the methodology, performed data preprocessing and analyses, developed the computational code, generated figures, and wrote the original draft of the manuscript.

André Mascioli Cravo contributed to statistical insights, methodological discussions, and interpretation of the results, and provided critical feedback throughout the analytical process.

Claudinei E. Biazoli Jr. supervised the entire research process, contributed to the conceptual development of the study, and critically revised and edited the manuscript.

All authors reviewed and approved the final version of the manuscript.

## 9. FUNDING

This study was financed by the CAPES (Coordenação de Aperfeiçoamento de Pessoal de Nível Superior – Brasil; grant n. 23006.027866/2022-27) and CNPq (INCT; National Institute of Science and Technology on Social and Affective Neuroscience, grant n. 406463/2022-0)

## 9 DECLARATION OF COMPETING INTERESTS

Tiago Duarte-Pereira: None; André Mascioli Cravo: None; Claudinei E. Biazoli Jr.: None

## Notes

### Competing Interest Statement

The authors have declared no competing interest.

## REFERENCES

Aliko, Sarah, Jiawen Huang, Florin Gheorghiu, Stefanie Meliss, and Jeremy I. Skipper. “A naturalistic neuroimaging database for understanding the brain using ecological stimuli”. Scientific Data 7, n°1 (13 de outubro de 2020): 347. 10.1038/s41597-020-00680-2.

Amico, Enrico, Francisco Gomez, Carol Di Perri, Audrey Vanhaudenhuyse, Damien Lesenfants, Pierre Boveroux, Vincent Bonhomme, Jean-François Brichant, Daniele Marinazzo, and Steven Laureys. “Posterior Cingulate Cortex-Related Co-Activation Patterns: A Resting State fMRI Study in Propofol-Induced Loss of Consciousness”. PLOS ONE 9, n° 6 (junho de 2014): 1–9. 10.1371/journal.pone.0100012.

Amico, Enrico, and Joaquín Goñi. “The Quest for Identifiability in Human Functional Connectomes”. Scientific Reports 8, n° 1 (29 de maio de 2018): 8254. 10.1038/s41598-018-25089-1.

Baek, Elisa C, Ryan Hyon, Karina López, Emily S. Finn, Mason A. Porter, and Carolyn Parkinson. “Popular Individuals Process the World in Particularly Normative Ways”, 4 de junho de 2021. 10.31234/osf.io/6fj2p.

Baldassano, Christopher, Janice Chen, Arianne Zadbood, Jonathan W. Pillow, Uri Hasson, and Kenneth A. Norman. “Discovering Event Structure in Continuous Narrative Perception and Memory”. Neuron 95, no. 3 (2017): 709–721. 10.1016/j.neuron.2017.06.041.

Barnes, Anna, Edward T. Bullmore, and John Suckling. “Endogenous Human Brain Dynamics Recover Slowly Following Cognitive Effort”. Organizado por Björn Brembs. PLoS ONE 4, n° 8 (14 de agosto de 2009): e6626. 10.1371/journal.pone.0006626.

Brunick, Kaitlin L., James E. Cutting, and Jordan E. DeLong. “Low-level features of film: What they are and why we would be lost without them.” Em Psychocinematics: Exploring cognition at the movies., 133–48. New York, NY, US: Oxford University Press, 2013. 10.1093/acprof:oso/9780199862139.003.0007.

Cabral, Joana, Morten L. Kringelbach, and Gustavo Deco. “Functional Connectivity Dynamically Evolves on Multiple Time-Scales over a Static Structural Connectome: Models and Mechanisms”. NeuroImage 160 (2017): 84–96. 10.1016/j.neuroimage.2017.03.045.

Cabral, Joana, Francesca Castaldo, Jakub Vohryzek, Vladimir Litvak, Christian Bick, Renaud Lambiotte, Karl Friston, Morten L. Kringelbach, and Gustavo Deco. “Metastable oscillatory modes emerge from synchronization in the brain spacetime connectome”. Communications Physics 5, n° 1 (15 de julho de 2022): 184. 10.1038/s42005-022-00950-y.

Cabral-Carvalho, Rodrigo M., Walter H. L. Pinaya, e João R. Sato. “A Graph Neural Network Approach to Investigate Brain Critical States Over Neurodevelopment”. Network Neuroscience (2025): 1–16. 10.1162/netn_a_00451.

Cavanna, Federico, Martina G. Vilas, Matías Palmucci, and Enzo Tagliazucchi. “Dynamic functional connectivity and brain metastability during altered states of consciousness”. NeuroImage 180 (2018): 383–95. 10.1016/j.neuroimage.2017.09.065.

Caviness, V. S. Jr, J. Meyer, N. Makris, and D. N. Kennedy. “MRI-Based Topographic Parcellation of Human Neocortex: An Anatomically Specified Method with Estimate of Reliability.” Journal of Cognitive Neuroscience 8, n° 6 (novembro de 1996): 566–87. 10.1162/jocn.1996.8.6.566.

Cutting, James E., Jordan E. DeLong, and Christine E. Nothelfer. “Attention and the Evolution of Hollywood Film”. Psychological Science 21, n°3 (2010): 432–39. 10.1177/0956797610361679.

Cohen, Samuel S., Nim Tottenham, and Christopher Baldassano. “Developmental Changes in Story-Evoked Responses in the Neocortex and Hippocampus”. ELife 11 (2022): e69430. 10.7554/eLife.69430.

Da Silva Castanheira, Jason, Hector D Orozco, Bratislav Misic, and Sylvain Baillet. “MEG, Myself, and I: Individual Identification from Neurophysiological Brain Activity”, 18 de fevereiro de 2021. 10.1101/2021.02.18.431803.

De Souza Rodrigues, Júlia, Fernanda Lenita Ribeiro, João Ricardo Sato, Rickson Coelho Mesquita, and Claudinei Eduardo Biazoli Júnior. “Identifying Individuals Using fNIRS-Based Cortical Connectomes”. Biomedical Optics Express 10, n° 6 (1° de junho de 2019): 2889. 10.1364/BOE.10.002889.

Dosenbach, Nico U. F., Marcus E. Raichle, and Evan M. Gordon. “The Brain’s Action-Mode Network”. Nature Reviews Neuroscience 26 (2025): 158–168. 10.1038/s41583-024-00895-x.

Fan, Siyan, Samaneh Nemati, Teddy J. Akiki, Jeremy Roscoe, Christopher L. Averill, Samar Fouda, Lynnette A. Averill, and Chadi G. Abdallah. “Pretreatment Brain Connectome Fingerprint Predicts Treatment Response in Major Depressive Disorder”. Chronic Stress 4 (2020): 2470547020984726. 10.1177/2470547020984726.

Finn, Emily S., Russell A. Poldrack, and James M. Shine. “Functional neuroimaging as a catalyst for integrated neuroscience”. Nature 623, n°7986 (1° de novembro de 2023): 263–73. 10.1038/s41586-023-06670-9.

Finn, Emily S., Xilin Shen, Dustin Scheinost, Monica D. Rosenberg, Jessica Huang, Marvin M. Chun, Xenophon Papademetris, and R. Todd Constable. “Functional Connectome Fingerprinting: Identifying Individuals Using Patterns of Brain Connectivity.” Nature Neuroscience 18, n°11 (novembro de 2015): 1664–71. 10.1038/nn.4135.

Fraiman, Daniel, Pablo Balenzuela, Jennifer Foss, and Dante R. Chialvo. “Ising-like dynamics in large-scale functional brain networks”. Phys. Rev. E 79, n° 6 (junho de 2009): 061922. 10.1103/PhysRevE.79.061922.

Frew, Simon, Ahmad Samara, Hallee Shearer, Jeffrey Eilbott, and Tamara Vanderwal. “Getting the nod: Pediatric head motion in a transdiagnostic sample during movie- and resting-state fMRI”. PLOS ONE 17, n°4 (abril de 2022): 1–19. 10.1371/journal.pone.0265112.

Glasser, Matthew, Timothy Coalson, Edward Robinson, et al. “A Multi-Modal Parcellation of Human Cerebral Cortex”. Nature 536 (2016): 171–178. 10.1038/nature18933.

Gonzalez-Castillo, Javier, Daniel A. Handwerker, Meghan E. Robinson, Colin Weir Hoy, Laura C. Buchanan, Ziad S. Saad, and Peter A. Bandettini. “The Spatial Structure of Resting State Connectivity Stability on the Scale of Minutes.” Frontiers in Neuroscience 8 (2014): 138. 10.3389/fnins.2014.00138.

Gordon, Evan M., Timothy O. Laumann, Babatunde Adeyemo, Jeremy F. Huckins, William M. Kelley, and Steven E. Petersen. “Generation and Evaluation of a Cortical Area Parcellation from Resting-State Correlations.” Cerebral Cortex (New York, N.Y.: 1991) 26, n° 1 (janeiro de 2016): 288–303. 10.1093/cercor/bhu239.

Gordon, Evan M., Timothy O. Laumann, Adrian W. Gilmore, Dillan J. Newbold, Deanna J. Greene, Jeffrey J. Berg, Mario Ortega, et al. “Precision Functional Mapping of Individual Human Brains.” Neuron 95, n° 4 (16 de agosto de 2017): 791-807.e7. 10.1016/j.neuron.2017.07.011.

Hancock, Fran, Fernando E. Rosas, Robert A. McCutcheon, Joana Cabral, Ottavia Dipasquale, and Federico E. Turkheimer. “Metastability as a candidate neuromechanistic biomarker of schizophrenia pathology”. PLOS ONE 18, n°3 (março de 2023): 1–35. 10.1371/journal.pone.0282707.

Hancock, F., Rosas, F.E., Luppi, A.I. et al. Metastability demystified — the foundational past, the pragmatic present and the promising future. Nat. Rev. Neurosci. 26, 82–100 (2025). 10.1038/s41583-024-00883-1.

Hanke, Michael, Nico Adelhöfer, Daniel Kottke, Vittorio Iacovella, Ayan Sengupta, Falko R. Kaule, Roland Nigbur, Alexander Q. Waite, Florian Baumgartner, and Jörg Stadler. “A studyforrest extension, simultaneous fMRI and eye gaze recordings during prolonged natural stimulation”. Scientific Data 3, n° 1 (25 de outubro de 2016): 160092. 10.1038/sdata.2016.92.

Hasson, Uri, Ohad Landesman, Barbara Knappmeyer, Ignacio Vallines, Nava Rubin, and David J. Heeger. “Neurocinematics: The Neuroscience of Film”. Projections 2, n° 1 (1° de janeiro de 2008): 1–26. 10.3167/proj.2008.020102.

Hasson, Uri, Yuval Nir, Ifat Levy, Galit Fuhrmann, and Rafael Malach. “Intersubject Synchronization of Cortical Activity During Natural Vision”. Science 303, n° 5664 (2004): 1634–40. 10.1126/science.1089506.

Hellyer, Peter J., Murray Shanahan, Gregory Scott, Richard J. S. Wise, David J. Sharp, and Robert Leech. “The Control of Global Brain Dynamics: Opposing Actions of Frontoparietal Control and Default Mode Networks on Attention.” The Journal of Neuroscience : The Official Journal of the Society for Neuroscience 34, n° 2 (8 de janeiro de 2014): 451–61. 10.1523/JNEUROSCI.1853-13.2014.

Herzog, Rubén, Pedro A. M. Mediano, Fernando E. Rosas, Paul Lodder, Robin Carhart-Harris, Yonatan Sanz Perl, Enzo Tagliazucchi, and Rodrigo Cofre. “A whole-brain model of the neural entropy increase elicited by psychedelic drugs”. Scientific Reports 13, n° 1 (17 de abril de 2023): 6244. 10.1038/s41598-023-32649-7.

Hudson, Michael, Kaisa Seppälä, Vesa Putkinen, Li Sun, Enrico Glerean, Tuomas Karjalainen, et al. “Dissociable Neural Systems for Unconditioned Acute and Sustained Fear”. NeuroImage 216 (2020): 116522. 10.1016/j.neuroimage.2020.116522.

Hutchison, R. Matthew, Melina Hutchison, Kathryn Y. Manning, Ravi S. Menon, and Stefan Everling. “Isoflurane Induces Dose-Dependent Alterations in the Cortical Connectivity Profiles and Dynamic Properties of the Brain’s Functional Architecture”. Human Brain Mapping 35, no. 12 (2014): 5754–5775. 10.1002/hbm.22583.

Hutchison, R. Matthew, Thilo Womelsdorf, Elena A. Allen, Peter A. Bandettini, Vince D. Calhoun, Maurizio Corbetta, Stefania Della Penna, et al. “Dynamic functional connectivity: Promise, issues, and interpretations”. NeuroImage 80 (2013): 360–78. 10.1016/j.neuroimage.2013.05.079.

Ibanez, Agustin, Morten L. Kringelbach, and Gustavo Deco. “A synergetic turn in cognitive neuroscience of brain diseases”. Trends in Cognitive Sciences, 2024. 10.1016/j.tics.2023.12.006.

Jangraw, David C., Emily S. Finn, Peter A. Bandettini, Nicole Landi, Haorui Sun, Fumiko Hoeft, Gang Chen, Kenneth R. Pugh, and Peter J. Molfese. “Inter-subject correlation during long narratives reveals widespread neural correlates of reading ability”. NeuroImage 282 (2023): 120390. 10.1016/j.neuroimage.2023.120390.

Kauttonen, Janne, Yevhen Hlushchuk, Iiro P. Jääskeläinen, and Pia Tikka. “Brain mechanisms underlying cue-based memorizing during free viewing of movie Memento”. NeuroImage 172 (2018): 313–25. 10.1016/j.neuroimage.2018.01.068.

Kringelbach, Morten L., Yonatan Sanz Perl, Enzo Tagliazucchi, and Gustavo Deco. “Toward naturalistic neuroscience: Mechanisms underlying the flattening of brain hierarchy in movie-watching compared to rest and task”. Science Advances 9, n° 2 (2023): eade6049. 10.1126/sciadv.ade6049.

Korponay, Christopher, Amanda C. Janes, and Benjamin B. Frederick. “Brain-Wide Functional Connectivity Artifactually Inflates throughout Functional Magnetic Resonance Imaging Scans”. Nature Human Behaviour 8, no. 8 (2024): 1568–1580. 10.1038/s41562-024-01908-6.

Lahnakoski, Juha M., Iiro P. Jääskeläinen, Mikko Sams, and Lauri Nummenmaa. “Neural mechanisms for integrating consecutive and interleaved natural events”. Human Brain Mapping 38, n° 7 (2017): 3360–76. 10.1002/hbm.23591.

Larabi, Daouia I., Martin Gell, Enrico Amico, Simon B. Eickhoff, and Kaustubh R. Patil. “Highly Accurate Local Functional Fingerprints and Their Stability”. bioRxiv 8, no. 03 (2021): 454862. 10.1101/2021.03.15.454862.

Lee, Christopher S., Mehraveh Aly, and Christopher Baldassano. “Anticipation of Temporally Structured Events in the Brain”. ELife 10 (2021): e64972. 10.7554/eLife.64972

Loschky, Lester C., Adam M. Larson, Joseph P. Magliano, and Tim J. Smith. “What Would Jaws Do? The Tyranny of Film and the Relationship between Gaze and Higher-Level Narrative Film Comprehension”. PLOS ONE 10, n° 11 (novembro de 2015): 1–23. 10.1371/journal.pone.0142474.

Maldjian, J. A., Laurienti, P. J., Kraft, R. A., & Burdette, J. H. (2003). An automated method for neuroanatomic and cytoarchitectonic atlas-based interrogation of fMRI data sets. Neuroimage, 19(3), 1233–1239. 10.1016/S1053-8119(03)00169-1.

Meer, Johan N. van der, Michael Breakspear, Luke J. Chang, Saurabh Sonkusare, and Luca Cocchi. “Movie viewing elicits rich and reliable brain state dynamics”. Nature Communications 11, n° 1 (5 de outubro de 2020): 5004. 10.1038/s41467-020-18717-w.

Nguyen, Vinh Thai, Saurabh Sonkusare, Jane Stadler, Xintao Hu, Michael Breakspear, and Christine Cong Guo. “Distinct Cerebellar Contributions to Cognitive-Perceptual Dynamics During Natural Viewing”. Cerebral Cortex 27, n° 12 (novembro de 2016): 5652–62. 10.1093/cercor/bhw334.

Ni, Yuqian, Xia Zheng, Richard Betzel, and Thomas W. James. “Increased Segregation in Functional Connectivity Networks When Watching Unpleasant Arousing Videos: A gPPI Analysis.” Brain Connectivity, 24 de janeiro de 2024. 10.1089/brain.2023.0048.

Parkinson, Carolyn, Adam M. Kleinbaum, and Thalia Wheatley. “Similar neural responses predict friendship”. Nature Communications 9, n°1 (30 de janeiro de 2018): 332. 10.1038/s41467-017-02722-7.

Raz, Gal, Yonatan Winetraub, Yael Jacob, Sivan Kinreich, Adi Maron-Katz, Galit Shaham, Ilana Podlipsky, Gadi Gilam, Eyal Soreq, and Talma Hendler. “Portraying emotions at their unfolding: A multilayered approach for probing dynamics of neural networks”. NeuroImage 60, n° 2 (2012): 1448–61. 10.1016/j.neuroimage.2011.12.084.

Reagan, Andrew J., Lewis Mitchell, Dilan Kiley, Christopher M. Danforth, and Peter Sheridan Dodds. “The emotional arcs of stories are dominated by six basic shapes”. EPJ Data Science 5, n°1 (4 de novembro de 2016): 31. 10.1140/epjds/s13688-016-0093-1.

Rieck, Bastian, Tristan Yates, Christian Bock, Karsten Borgwardt, Guy Wolf, Nicholas Turk-Browne, and Smita Krishnaswamy. “Uncovering the topology of time-varying fMRI data using cubical persistence”. Em Proceedings of the 34th International Conference on Neural Information Processing Systems. NIPS’20. Red Hook, NY, USA: Curran Associates Inc., 2020.

Saarimäki, Heini. “Naturalistic Stimuli in Affective Neuroimaging: A Review”. Frontiers in Human Neuroscience 15 (2021). 10.3389/fnhum.2021.675068.

Sadeghi, Fatemeh, Elvira del Agua Banyeres, Alessandra Pizzuti, Abdullah Okar, Kai Grimm, Christian Gerloff, Morten L. Kringelbach, Rainer Goebel, Simone Zittel, and Gustavo Deco. “The Arrow of Time in Parkinson’s Disease”. NeuroImage: Clinical 47 (2025): 103834. 10.1016/j.nicl.2025.103834.

Sato, João Ricardo, Claudinei Eduardo Biazoli, André Zugman, Pedro Mario Pan, Ana Paula Arantes Bueno, Luciana Monteiro Moura, Ary Gadelha, et al. “Long-term Stability of the Cortical Volumetric Profile and the Functional Human Connectome throughout Childhood and Adolescence”. European Journal of Neuroscience 54, n° 6 (setembro de 2021): 6187–6201. 10.1111/ejn.15435.

Sonkusare, Saurabh, Michael Breakspear, and Christine Guo. “Naturalistic Stimuli in Neuroscience: Critically Acclaimed”. Trends in Cognitive Sciences 23, n° 8 (2019): 699–714. 10.1016/j.tics.2019.05.004.

Sorrentino, Pierpaolo, Rosaria Rucco, Anna Lardone, Marianna Liparoti, Emahnuel Troisi Lopez, Carlo Cavaliere, Andrea Soricelli, Viktor Jirsa, Giuseppe Sorrentino, and Enrico Amico. “Clinical connectome fingerprints of cognitive decline”. NeuroImage 238 (2021): 118253. 10.1016/j.neuroimage.2021.118253.

Sorrentino, Pierpaolo, Emahnuel Troisi Lopez, Antonella Romano, Carmine Granata, Marie Constance Corsi, Giuseppe Sorrentino, and Viktor Jirsa. “Brain fingerprint is based on the aperiodic, scale-free, neuronal activity”. NeuroImage 277 (15 de agosto de 2023): 120260. 10.1016/j.neuroimage.2023.120260.

Tagliazucchi, Enzo, Robin Carhart-Harris, Robert Leech, David Nutt, and Dante R. Chialvo. “Enhanced Repertoire of Brain Dynamical States during the Psychedelic Experience”. Human Brain Mapping 35, no. 11 (2014): 5442–5456. 10.1002/hbm.22562.

Tagliazucchi, Enzo, and Helmut Laufs. “Decoding Wakefulness Levels from Typical fMRI Resting-State Data Reveals Reliable Drifts between Wakefulness and Sleep.” Neuron 82, n° 3 (7 de maio de 2014): 695–708. 10.1016/j.neuron.2014.03.020.

Tepper, Ángeles, Javiera Vásquez Núñez, Juan Pablo Ramirez-Mahaluf, Juan Manuel Aguirre, Daniella Barbagelata, Elisa Maldonado, Camila Díaz Dellarossa, et al. “Intra and inter-individual variability in functional connectomes of patients with First Episode of Psychosis”. NeuroImage: Clinical 38 (2023): 103391. 10.1016/j.nicl.2023.103391.

Tong, Yunjie, Laura M. Hocke, and Benjamin B. Frederick. “Low Frequency Systemic Hemodynamic ‘Noise’ in Resting State BOLD fMRI: Characteristics, Causes, Implications, Mitigation Strategies, and Applications”. Frontiers in Neuroscience 13 (2019): 787. 10.3389/fnins.2019.00787.

Vanderwal, Tamara, Jeffrey Eilbott, Clare Kelly, Simon R. Frew, Todd S. Woodward, Michael P. Milham, and F. Xavier Castellanos. “Stability and Similarity of the Pediatric Connectome as Developmental Measures.” NeuroImage 226 (1° de fevereiro de 2021): 117537. 10.1016/j.neuroimage.2020.117537.

Vanderwal, Tamara, Clare Kelly, Jeffrey Eilbott, Linda C. Mayes, and F. Xavier Castellanos. “Inscapes: A Movie Paradigm to Improve Compliance in Functional Magnetic Resonance Imaging.” NeuroImage 122 (15 de novembro de 2015): 222–32. 10.1016/j.neuroimage.2015.07.069.

Ville, Dimitri Van De, Younes Farouj, Maria Giulia Preti, Raphaël Liégeois, and Enrico Amico. “When makes you unique: Temporality of the human brain fingerprint”. Science Advances 7, n° 42 (2021): 10.1126/sciadv.abj0751.

Wallace, Raven S., Brontë Mckeown, Ian Goodall-Halliwell, Louis Chitiz, Philippe Forest, Theodoros Karapanagiotidis, Bridget Mulholland, et al. “Mapping Patterns of Thought onto Brain Activity during Movie-Watching”, 31 de janeiro de 2024. 10.1101/2024.01.31.578244.

Wang, Jiahui, Yudan Ren, Xintao Hu, Vinh Thai Nguyen, Lei Guo, Junwei Han, e Christine Cong Guo. “Test–retest reliability of functional connectivity networks during naturalistic fMRI paradigms”. Human Brain Mapping 38, n° 4 (2017): 2226–41. 10.1002/hbm.23517.

Wens, Vincent, Mathieu Bourguignon, Marc Vander Ghinst, Alison Mary, Brice Marty, Nicolas Coquelet, Gilles Naeije, Philippe Peigneux, Serge Goldman, e Xavier De Tiège. “Synchrony, metastability, dynamic integration, and competition in the spontaneous functional connectivity of the human brain”. NeuroImage 199 (2019): 313–24. 10.1016/j.neuroimage.2019.05.081.

