## Supplementary Material for "Connectome Similarity Varies with Global Metastability and Movie Content"

### Additional descriptive statistics

| | **Iself** | | |
|---|---|---|---|
| **Movies** | **30s** | **300s** | **600s** |
| 12 Years a Slave | M=0.260,SE=0.002, CI[0.182 0.333] | M=0.560,SE=0.005, CI[0.492 0.624] | M=0.694,SE=0.006, CI[0.642 0.747] |
| 500 Days of Summer | M=0243, SE=0.001, CI[0.176 0.310] | M=0.533, SE=0.003, CI[0.465 0.601] | M=0.658, SE=0.004, CI[0.597 .718] |
| Back to the Future | M=0.236, SE=0.001, CI[0.168 0.304] | M=0.544, SE=0.005, CI[0.483 0.606] | M=0.688, SE=0.006, CI[0.640 0.737] |
| Citizenfour | M=0.247, SE=0.001, CI[0.173 0.320] | M=0.545, SE=0.003, CI[0.475 0.615] | M=0.676, SE=0.004, CI[0.616 0.736] |
| Little Miss Sunshine | M=0.250, SE=0.003, CI[0.179 0.321] | M=0.554, SE=0.006, 0.487 0.620] | M=0.684, SE=0.008, CI[0.624 0.743] |
| Pulp Fiction | M=0.247, SE=0.002, CI[0.169 0.324] | M=0.532, SE=0.006, CI[0.453 0.611] | M=0.670, SE=0.007, CI[0.606 0.732] |
| Split | M=0.245, SE=0.002, CI[0.182 0.307] | M=0.545, SE=0.004, CI[0.498 0.624] | M=0.677, SE=0.005, CI[0.636 0.719] |
| The Prestige | M=0.250, SE=0.002, CI[0.178 0.320] | M=0.561, SE=0.005, CI[0.498 0.624] | M=0.703, SE=0.005, CI[0.658 0.748] |
| The Shawshank Redemption | M=0.246, SE=0.002, CI[0.180 0.312] | M=0.545, SE=0.004, CI[0.487 0.604] | M=0.681, SE=0.004, CI[0.646 0.717] |
| Usual Suspects | M=0.234, SE=0.002, CI[0.171 0.298] | M=0.523, SE=0.005, CI[0.460 0.587] | M=0.644, SE=0.006, CI[0.595 0.693] |

| | **Iothers** | | |
|---|---|---|---|
| **Movies** | **30s** | **300s** | **600s** |
| 12 Years a Slave | M=0.120, SE=0.0005, CI[0.104 0.135] | M=0.302, SE=0.001, CI[0.285 0.319] | M=0.384, SE=0.002, CI[0.367 0.401] |
| 500 Days of Summer | M=0.113, SE=0.0003, CI[0.098 0.128] | M=0.296, SE=0.001, CI[0.274 0.318] | M=0.381, SE=0.001, CI[0.357 0.405] |
| Back to the Future | M=0.111, SE=0.0006, CI[0.095 0.127] | M=0.291, SE=0.001, CI[0.271 0.311] | M=0.373, SE=0.002, CI[0.351 0.395] |
| Citizenfour | M=0.114, SE=0.0003, CI[0.100 0.128] | M=0.300, SE=0.001, CI[0.280 0.312] | M=0.384, SE=0.001, CI[0.361 0.408] |

| | | | |
|---|---|---|---|
| Little Miss Sunshine | M=0.115, SE=0.0006, CI[0.098 0.131] | M=0.294, SE=0.002, CI[0.273 0.316] | M=0.377, SE=0.003, CI[0.356 0.400] |
| Pulp Fiction | M=0.115, SE=0.0005, CI[0.098 0.132] | M=0.303, SE=0.001, CI[0.281 0.325] | M=0.388, SE=0.002, CI[0.365 0.411] |
| Split | M=0.132, SE=0.0009, CI[0.108 0.155] | M=0.340, SE=0.002, CI[0.317 0.363] | M=0.431, SE=0.002, CI[0.411 0.450] |
| The Prestige | M=0.117, SE=0.0006, CI[0.100 0.134] | M=0.305, SE=0.002, CI[0.283 0.326] | M=0.386, SE=0.002, CI[0.362 0.411] |
| The Shawshank Redemption | M=0.117, SE=0.0006, CI[0.098 0.131] | M=0.305, SE=0.002, CI[0.280 0.331] | M=0.390, SE=0.002, CI[0.362 0.417] |
| Usual Suspects | M=0.115, SE=0.0006, CI[0.098 0.131] | M=0.308, SE=0.001, CI[0.288 0.328] | M=0.398, SE=0.002, CI[0.380 0.418] |

| | **Metastability** | | |
|---|---|---|---|
| **Movies** | **30s** | **300s** | **600s** |
| 12 Years a Slave | M= 0.098, SE=0.002, CI[0.044 0.152] | M=0.110, SE=0.003, CI[0.067 0.152] | M=0.111, SE=0.046, CI[0.072 0.150] |
| 500 Days of Summer | M=0.088, SE=0.001, CI[0.040 0.136] | M=0.099, SE=0.002, CI[0.057 0.141] | M=0.101, SE=0.003, CI[0.062 0.141] |
| Back to the Future | M=0.090, SE=0.002, CI[0.042 0.138] | M=0.103, SE=0.004, CI[0.058 0.149] | M=0.106, SE=0.005, CI[0.062 0.150] |
| Citizenfour | M=0.089, SE=0.001, CI[0.048 0.129] | M=0.100, SE=0.001, CI[0.068 0.133] | M=0.103, SE=0.002, CI[0.072 0.133] |
| Little Miss Sunshine | M=0.092, SE=0.001, CI[0.054 0.130] | M=0.103, SE=0.003, CI[0.070 0.136] | M=0.104, SE=0.004, CI[0.071 0.137] |
| Pulp Fiction | M=0.077, SE=0.001, CI[0.043 0.110] | M=0.086, SE=0.002, CI[0.053 0.118] | M=0.088, SE=0.003, CI[0.057 0.118] |
| Split | M=0.086, SE=0.015, CI[0.047 0.125] | M=0.095, SE=0.002, CI[0.062 0.128] | M=0.097, SE=0.003, CI[0.066 0.128] |
| The Prestige | M=0.099, SE=0.001, CI[0.050 0.147] | M=0.111, SE=0.003, CI[0.075 0.147] | M=0.112, SE=0.003, CI[0.080 0.144] |
| The Shawshank Redemption | M=0.083, SE=0.011, CI[0.051 0.115] | M=0.092, SE=0.002, CI[0.066 0.117] | M=0.093, SE=0.002, CI[0.070 0.117] |
| Usual Suspects | M=0.084, SE=0.001, CI[0.044 0.125] | M=0.098, SE=0.003, CI[0.064 0.132] | M=0.101, SE=0.004, CI[0.071 0.132] |

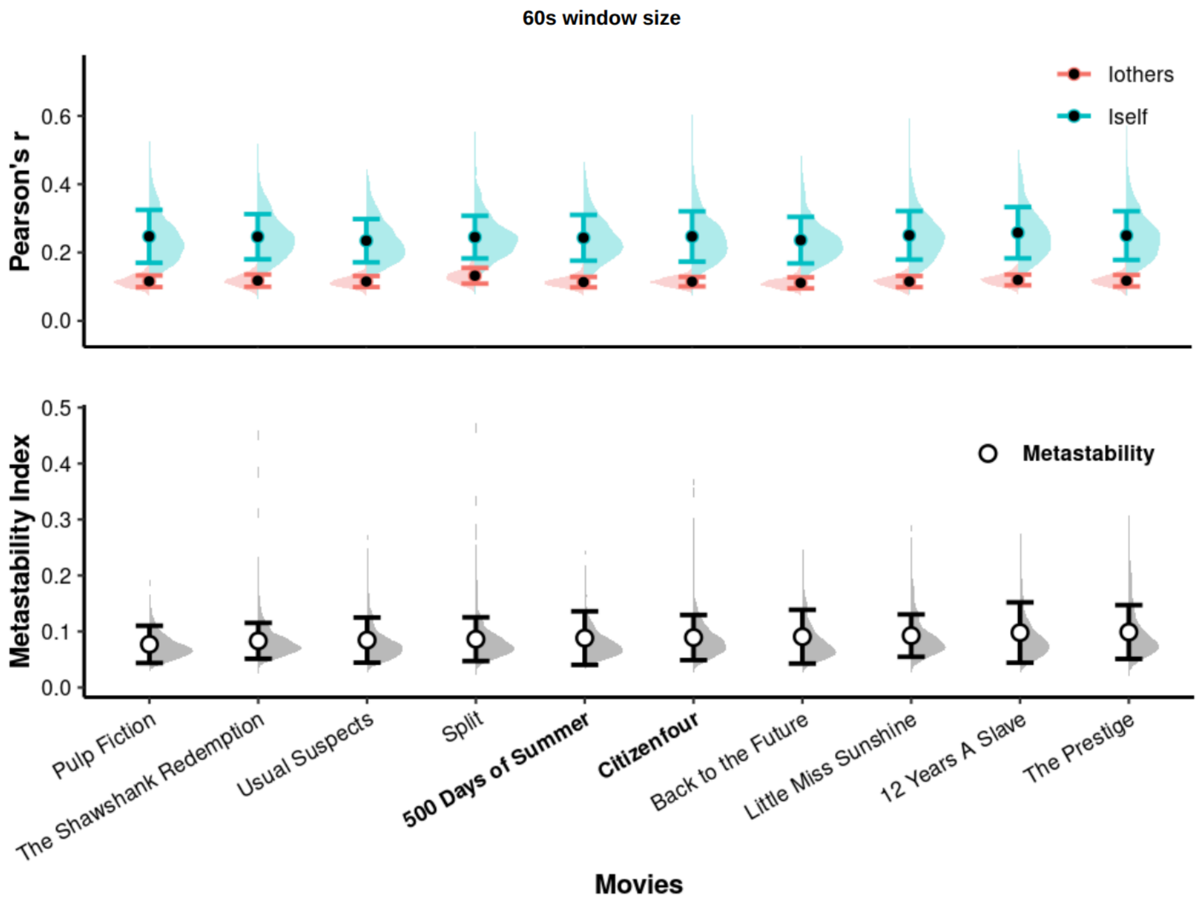
60s window size
Pearson's r
0.6
0.4
0.2
0.0
Iothers
Iself
Metastability Index
0.5
0.4
0.3
0.2
0.1
0.0
Metastability
Pulp Fiction
The Shawshank Redemption
Usual Suspects
Split
500 Days of Summer
Citizenfour
Back to the Future
Little Miss Sunshine
12 Years A Slave
The Prestige
Movies

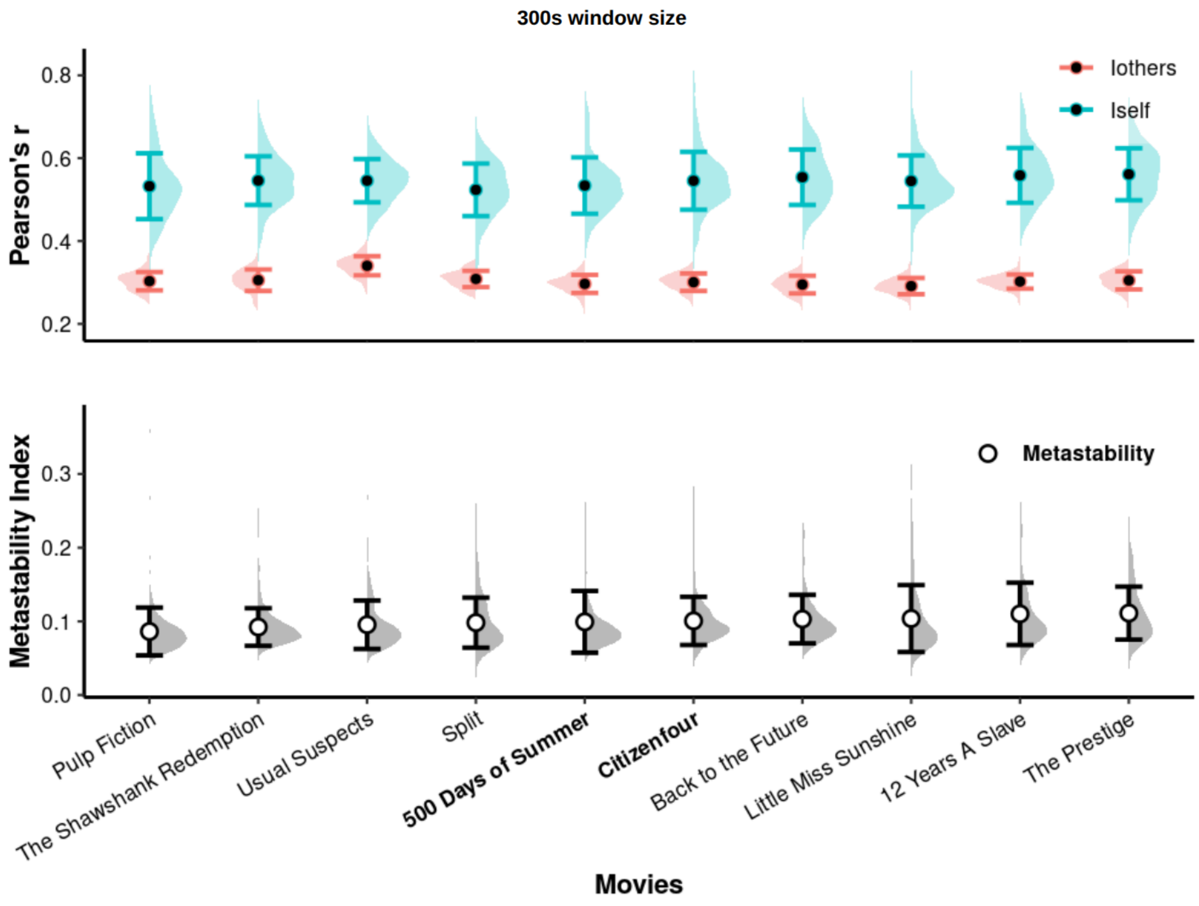


### Iself, Iothers and Metastability - Two movies - Across data (60-seconds)

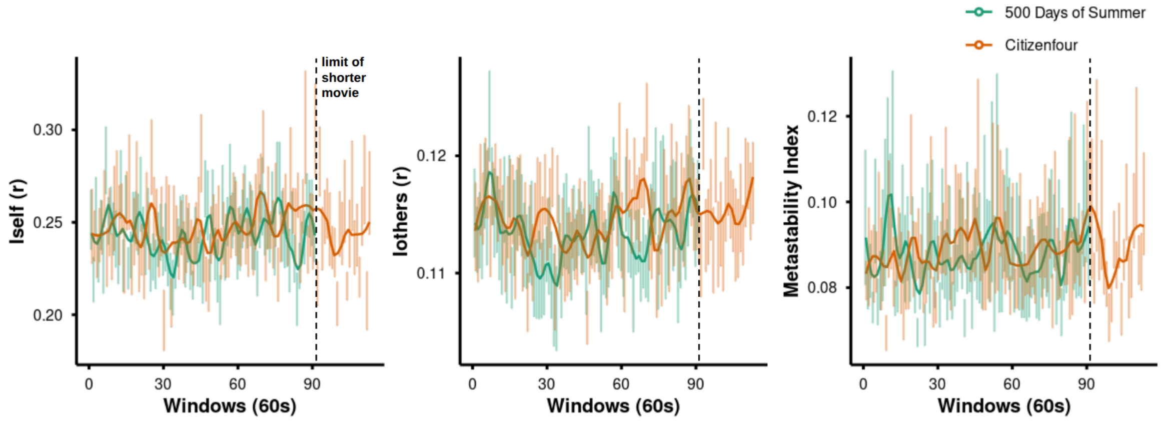


### Iself, Iothers and Metastability - Two movies - Across data (300-seconds)

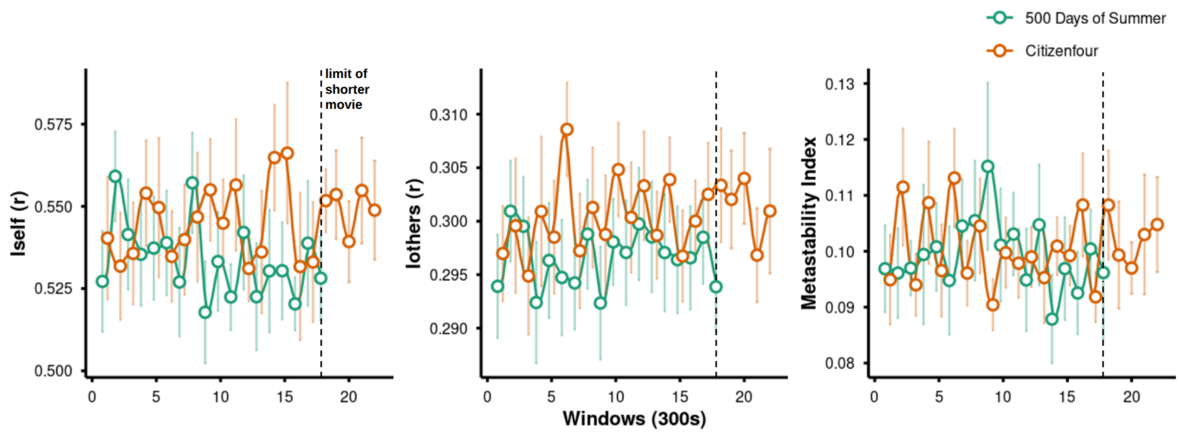


**S1 - Does differences between window sizes explain the association between metastability and identifiability metrics?**

Below, we present the other window sizes predictions for both indexes ranging from 120-seconds to 540-seconds, demonstrating the same effect and tendencies of the reported window sizes.

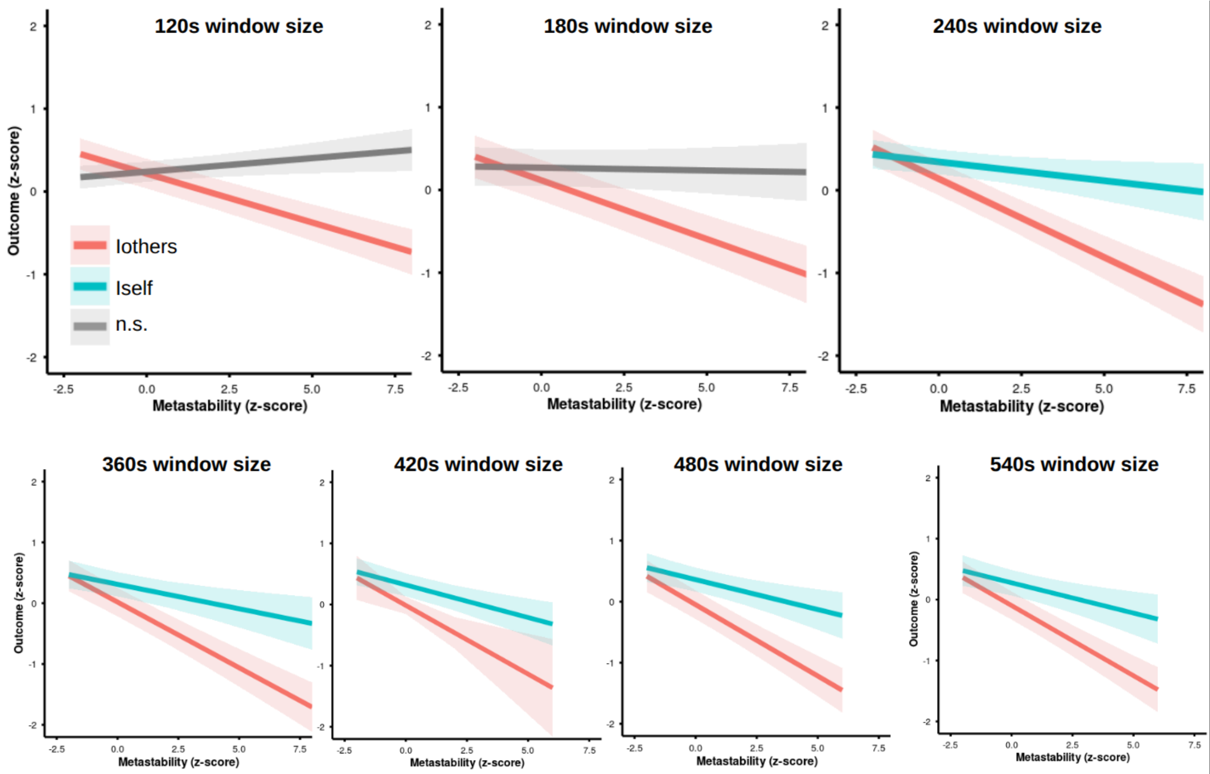

**S2 - Does differences between parcelations explain the association between metastability and identifiability metrics?**

All three atlases presented stability on the metrics calculation across window sizes. As already expected, the Harvard-Oxford atlas showed some differences compared to the other two, probably due to less parcels. It had a considering lower fingerprinting percentage for small windows and a still slight difference as more information is considered into the window. (Figure 3.A). Also, correlating average Iself and Iothers time series across atlases and windows, a stronger correlation between Glasser and Gordon atlas its visible in both metrics; in the other hand, Harvard-Oxford atlas correlations showed a consistently weaker correlation, but still satisfactory, except for a bigger window size like 600 seconds, which is also reasonable considering less timepoints for correlation. (Figure 3.B). And the average global metastability across atlases and window sizes also doesn't present any significant difference. (Figure 3.C).

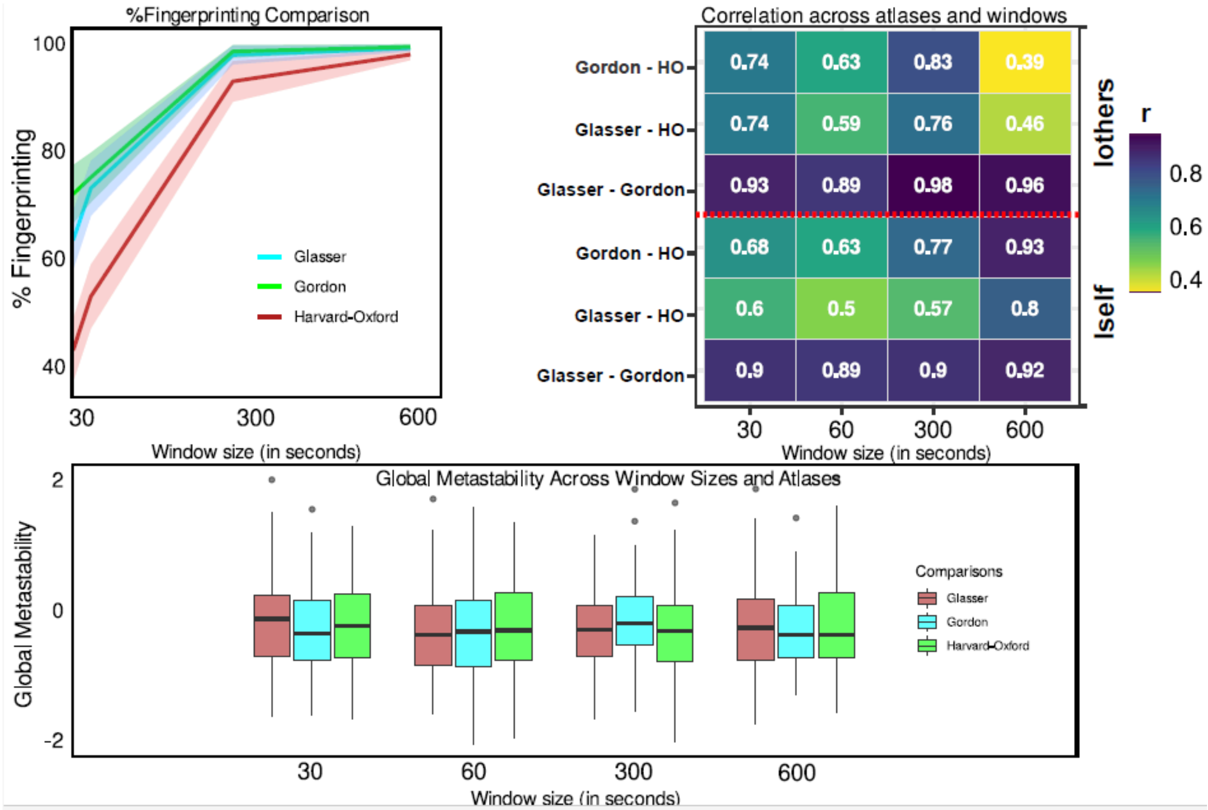

In the model, the association between Global Metastability and identification metrics (Iothers and Iself) was found across windows independently of the atlases. For this model, Iothers drives the most part on the prediction, and Iself and Windows as complementary predictors. As before, Harvard-Oxford was the worst for modelling the association, but even losing statistical significance in wider windows (300 seconds and 600 seconds), it follows the same pattern which in Glasser and Gordon atlas is negatively enhanced as more information gets into the window. These results clarify that the association between nonlinear dynamics and identification in naturalistic fMRI data isn't an atlas artifact, and the rest of investigation can move on focusing on analysis made with an adequate atlas. For this work, further investigation will be made based on Gordon atlas, for presenting adequate performance for modelling the association. (Figure 4).

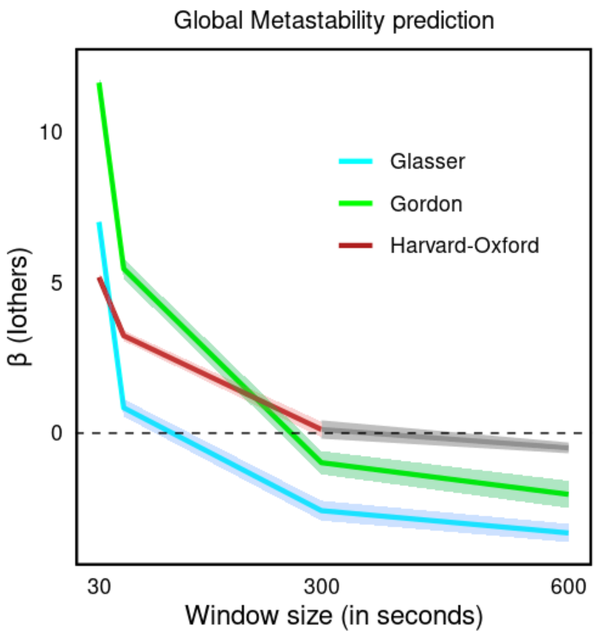


**S3 - Does sLFO artifacts significantly affect the data?**

Scans with longer than usual duration, such as those during whole-movies watching, are potentially confounded by systemic low-frequency oscillations (sLFO) artifacts. These artifacts cause considerable FC inflation. In a recent paper, Korponay et al. (2024) shed light to how these artifacts usually persist after standard preprocessing for longer scans. As expected, we observed this sLFO artifact in the NNdb dataset, but with a stabilization of FC measures after 16 minutes (Figure 5).

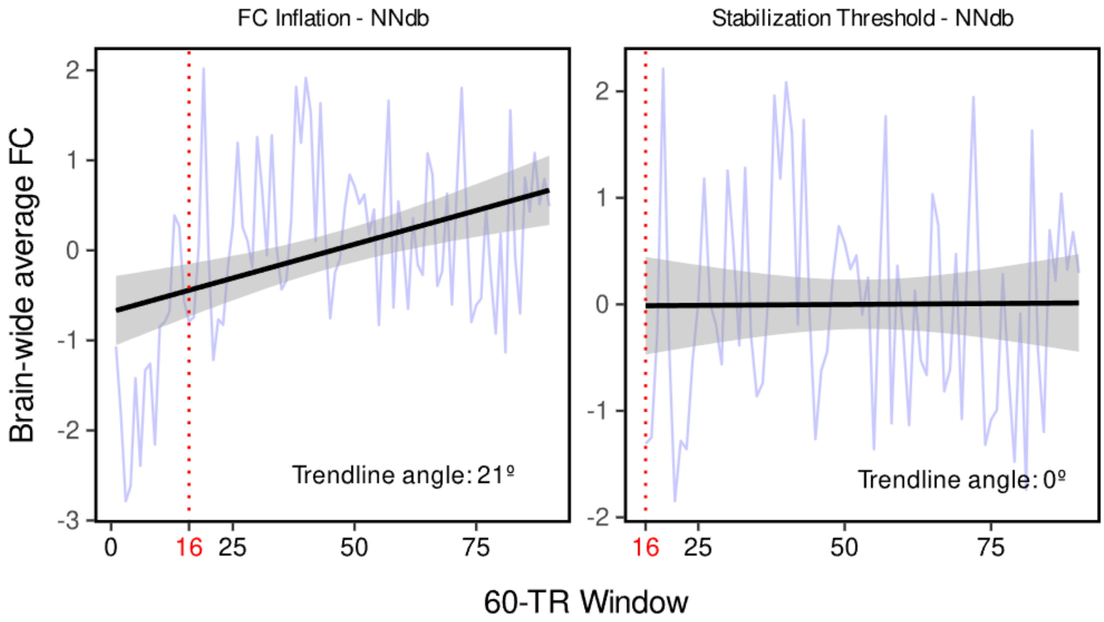


**Figure 5:** The FC inflation (left image) based on sLFO was indeed found in the NNDb (Neuroimaging Naturalistic Database), showing a positive trend of 21° across the time series but restricted to the initial minutes of the data, stabilizing after 16 minutes (right image). Considering the nature and structure of our analysis for the entire movie length, we chose to work with the data in raw state. Future studies should pay closer attention to the nature of sLFO.

At a movie-level analysis, no evidence for FC inflation was observed for *Citizenfour.* For *500 Days of Summer*, we compare the levels of fingerprinting percentage, Iself, Iothers and Global Metastability (i.e. spatial coherence through time) into 4 different window sizes (30, 60, 300 and 600 seconds) for the 'No Filter' and 'With Filter' data (Figure 6). No significant difference between the filtered and the no filtered data was found for the fingerprinting and global metastability measures. Based on these results, for the aim of this work, using a sLFO filter doesn't seem to be necessary. Further investigations should be conducted to refine its use on long-naturalistic datasets and to clarify how arousal-based inflation is related with the content of movies.

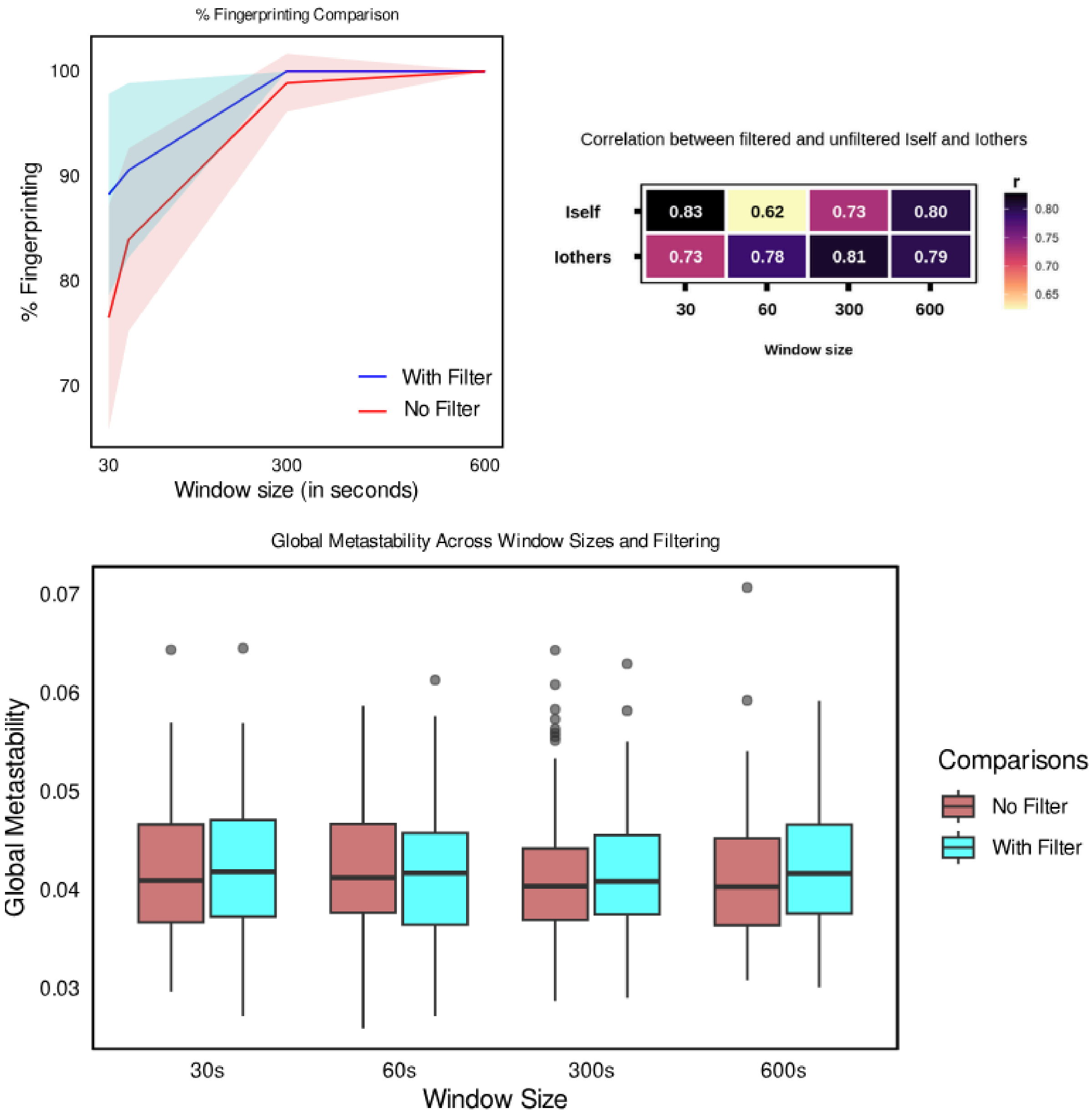


**Figure 6: Upper left:** A slightly non-significant higher percentage of fingerprinting for this dataset was found for the filtered version in narrow windows; in wider windows this gap tends to fade. **Upper right:** The correlation between the average Iself and Iothers, filtered and unfiltered, maintained a high positive correlation through the windows. **Bottom:** No significant difference was found for the average global metastability compared with the filtered version through the window sizes.

### S4 - Does movement parameters explain the association between metastability and identifiability metrics?

One common hypothesis in neuroimaging is that the results found are based on movement artifacts. To show that isn't the case, we extract the six realignment parameters (Left and Right; Anterior and Posterior; Inferior and Superior; and rotational component for each axes: Rotation X; Rotation Y; and Rotation Z) calculated on Conn Toolbox for each subject across all time-windows predicting global metastability; for reference example, we present these results for the 600s time-window below, showing that there's no such relationship between global metastability and movement parameters.

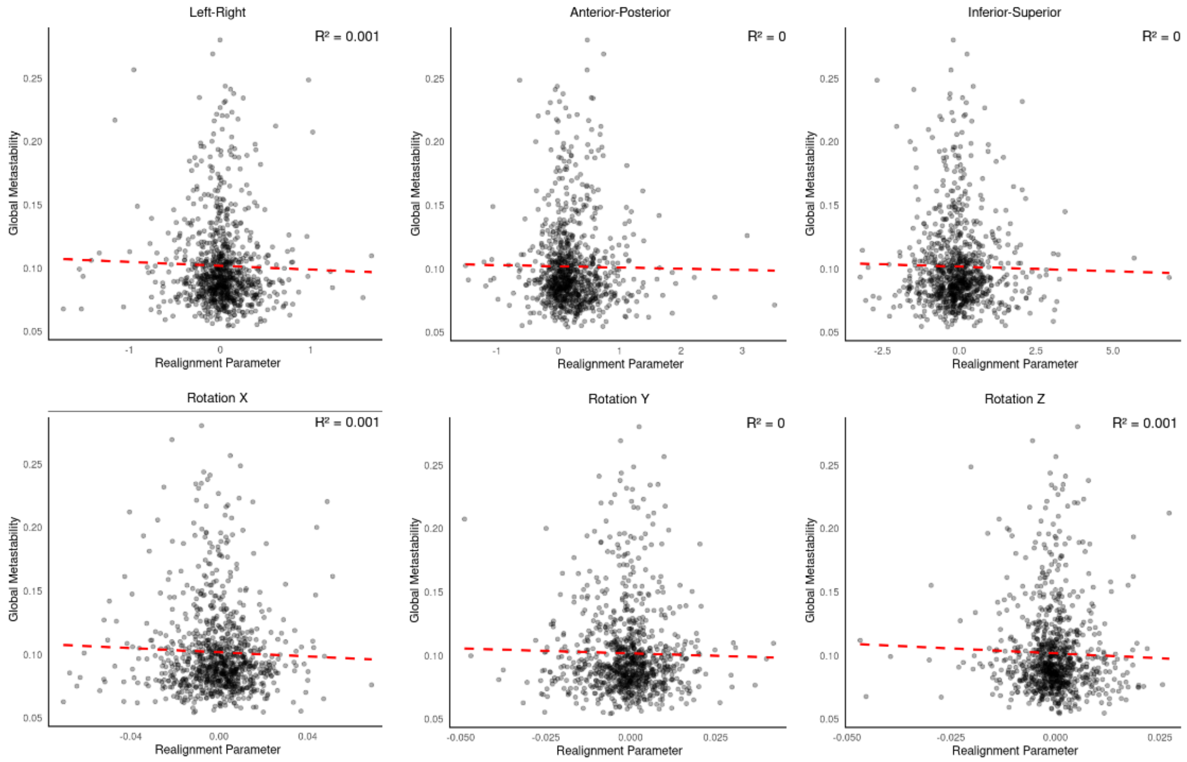


**Figure 7:** Prediction of global metastability by six motion parameters axes: Left and Right; Anterior and Posterior; Inferior and Superior; Rotation X; Rotation Y; Rotation Z. Considering the 600-seconds time-window and all subjects (N=86) data, no significant prediction was computed in any axes,; and presented flatter slopes (red dotted line).

### Additional Discussion

While nonlinear brain dynamics can be readily characterized by avalanches patterns in MEG, metastability is a suitable and reliable measurement of such dynamics in fMRI (Hellyer et al., 2014; Hancock et al., 2025). We found mostly consistent associations between metastability and fingerprinting for different time resolutions, spatial scales and stimuli contents. On the other hand, some differences in the associations between fingerprinting indexes and metastability were observed, in line with the expected non-linear brain dynamics (Cabral et al., 2017). For time scales in particular, using fixed time windows may introduce artifacts in our analyses, particularly for movie content. A natural extension would include implementing data-driven event segmentation to define proper time windows (Baldassano et al., 2017).
